# Brain-wide neural recordings in mice navigating physical spaces enabled by a cranial exoskeleton

**DOI:** 10.1101/2023.06.04.543578

**Authors:** James Hope, Travis Beckerle, Pin-Hao Cheng, Zoey Viavattine, Michael Feldkamp, Skylar Fausner, Kapil Saxena, Eunsong Ko, Ihor Hryb, Russell Carter, Timothy Ebner, Suhasa Kodandaramaiah

## Abstract

Complex behaviors are mediated by neural computations occurring throughout the brain. In recent years, tremendous progress has been made in developing technologies that can record neural activity at cellular resolution at multiple spatial and temporal scales. However, these technologies are primarily designed for studying the mammalian brain during head fixation – wherein the behavior of the animal is highly constrained. Miniaturized devices for studying neural activity in freely behaving animals are largely confined to recording from small brain regions owing to performance limitations. We present a cranial exoskeleton that assists mice in maneuvering neural recording headstages that are orders of magnitude larger and heavier than the mice, while they navigate physical behavioral environments. Force sensors embedded within the headstage are used to detect the mouse’s milli-Newton scale cranial forces which then control the x, y, and yaw motion of the exoskeleton via an admittance controller. We discovered optimal controller tuning parameters that enable mice to locomote at physiologically realistic velocities and accelerations while maintaining natural walking gait. Mice maneuvering headstages weighing up to 1.5 kg can make turns, navigate 2D arenas, and perform a navigational decision-making task with the same performance as when freely behaving. We designed an imaging headstage and an electrophysiology headstage for the cranial exoskeleton to record brain-wide neural activity in mice navigating 2D arenas. The imaging headstage enabled recordings of Ca^2+^ activity of 1000s of neurons distributed across the dorsal cortex. The electrophysiology headstage supported independent control of up to 4 silicon probes, enabling simultaneous recordings from 100s of neurons across multiple brain regions and multiple days. Cranial exoskeletons provide flexible platforms for largescale neural recording during the exploration of physical spaces, a critical new paradigm for unraveling the brain-wide neural mechanisms that control complex behavior.

## INTRODUCTION

A central goal in neuroscience is to understand how the brain mediates behavior. While traditionally neuroscientists studied each distinct brain region in isolation to determine its contributions to behavior, recent advances in recording technologies now allow neuroscientists to study the simultaneous activity of 1000s of neurons distributed across the brain. Advanced technologies such as high-density silicon neural recording probes (Neuropixels)^1–4^ and mesoscale optical imaging systems^5–9^ have revealed that neural substrates of behavior are distributed across multiple anatomically and functionally distinct brain regions. Limiting the use of these technologies are both the large form factor and weight, which can be orders of magnitude heavier than the rodent subjects. For instance, mice, the most commonly used animal model in neuroscience, typically weigh 20-40 g.

Most large-scale recordings require immobilizing the mice using head restraints, significantly limiting the behavioral repertoire. To overcome some limitations of head-fixation, immersive virtual reality environments^10, 11^, voluntary head-fixation of mice^12^ and rats^13^, floating environments^14, 15^, and rotating head-restraint^10^ have been developed. However, the lack of vestibular inputs that provide the sense of motion, balance, and orientation^16^, disruption in eye-head movement coupling^17^, and behavioral effects from increased stress can significantly alter neural activity during head-fixation compared to freely behaving^18^. While several head-mounted, miniaturized devices have been developed for imaging and recording neural activity in freely moving animals^9, 19–21^, these devices are typically limited to ∼10% (2-4 g) of a mouse’s body weight, resulting in sharp tradeoffs in device performance and capabilities.

Here we take a fundamentally different approach that eliminates the need for drastic miniaturization of neural interfaces: a cranial exoskeleton that uses force feedback from the mouse’s head to actuate a headstage that is orders of magnitude heavier and larger than a typical mouse (mouse: ∼3x10^-2^ kg, 30 cm^3^; headstage: 1.5 kg, 3 x10^3^ cm^3^; exoskeleton: 30 kg, 1x10^6^ cm^3^; **Fig. 1a-b**). We discovered controller tuning parameters that enabled mice to walk with velocities and accelerations comparable to freely behaving mice in open field arenas while maintaining natural gait. Mice maneuvering the exoskeleton-actuated headstages learned to locomote in 2D behavioral arenas and make sensory cue-guided navigational decisions with high proficiency (**Fig. 1d-e**). We developed two versions of exoskeleton-actuated headstage: one for imaging and one for electrophysiology. The imaging headstage enabled the recording of single cell activities from 1000s of cells distributed throughout the dorsal cortex as mice navigated a 2D arena (**Fig. 1e-f**). The electrophysiology headstage was capable of simultaneously lowering up to 4 neural probes into the brain and enabled multi-site, multi-day recordings from 261 neurons in 6 different brain regions during a navigational decision-making task (**Fig. 1g-h**).

**Fig. 1:**
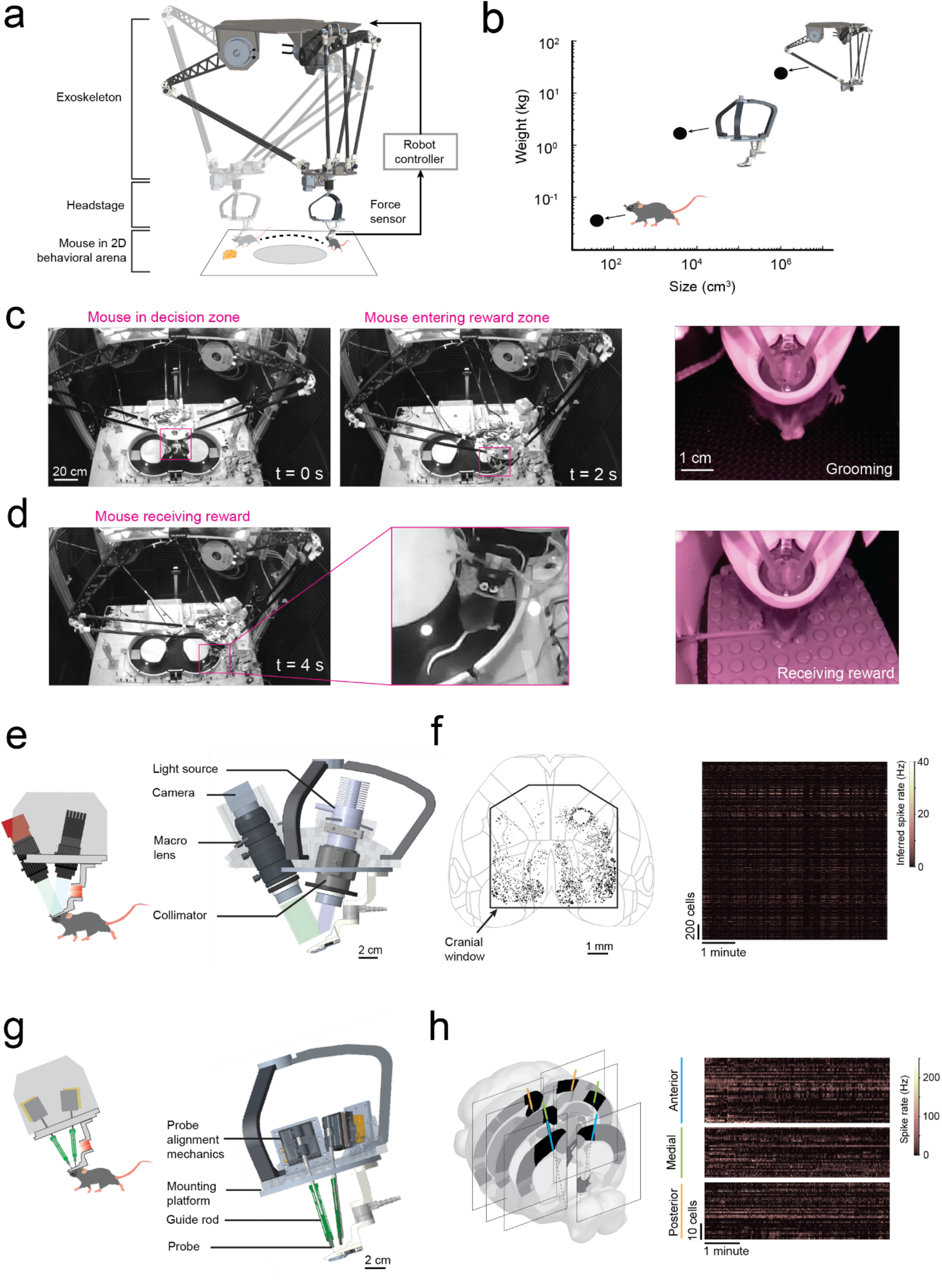
Cranial exoskeleton assisted brain wide neural recordings in mice navigating physical spaces (a) Schematic of the overall system: (top) robotic exoskeleton, (right) force feedback controller, (middle) headstage for mounting neural and behavioral recording hardware, (bottom) mouse in behavioral arena. (b) Log scale plot of size versus mass with points indicating a mouse, headstage, and exoskeleton. (c) System camera view of a mouse maneuvering the exoskeleton navigating a 2D behavioral arena (pink box around mouse), and (right) headstage-mounted camera view of a mouse grooming. (d) System camera view and close-up of a mouse maneuvering the exoskeleton receiving a reward, and (right) headstage-mounted camera view of the same event. (e) CAD rendering of headstage for mesoscale imaging at cellular resolution. (f) Locations of all cells imaged using the headstage for mesoscale imaging from a recording session of one mouse navigating a 2D arena, and (right) the inferred spike rates of the cells. (g) CAD rendering of headstage for electrophysiology. (h) 6 probe insertion sites recorded from across 3 sessions in a mouse performing a navigational decision-making task in a 2D maze, and (right) the spike rates recorded cells.

## RESULTS

### Rodent cranial exoskeleton design and construction

The cranial exoskeleton consisted of a 3-armed parallel robot, called a delta robot, to generate translational motion in the x, y, and z axes, and a motorized goniometer mounted to the moving platform of the delta robot to generate rotational motion in the pitch, roll, and yaw axes (**Supplementary Info. 1a-c**). The distal end of the goniometer was coupled to the top of the headstage, and a slipring was placed around this coupling to provide electrical access to equipment on the headstage (**Supplementary Info. 1d**). Mice were docked to the bottom of the headstage using a kinematic clamping mechanism that attached to a chronically implanted titanium headpost (**Supplementary Info. 1e**). The mice controlled their velocity and acceleration by applying forces through this clamping mechanism. These forces were measured using a 6-axis, milli-Newton scale force sensor embedded within the headstage close to the mouse, and then passed onto a force feedback controller that used an admittance control law to compute the desired velocity and acceleration. We sought to design the exoskeleton to be capable of moving at up to 0.2 m/s velocity and 1 m/s^2^ acceleration in the linear axes, and 180 deg/s velocity and 540 deg/s^2^ acceleration in the rotational axes based on previous studies characterizing unrestrained mouse behavior using markerless tracking^9^ and head-borne inertial measurement units^17, 22^. We set the desired range of motion to 80 x 80 cm in x and y to encompass an open field behavioral arena, with infinite rotation in yaw (**Supplementary Info. 4a-c**).

The primary consideration when designing the exoskeleton is that the linkage dimensions and motor torques must be capable of maneuvering the headstage at physiologically realistic velocities and accelerations across the full span of the behavioral arena. To evaluate this, we performed *in silico* simulations on kinematic and dynamic models of the exoskeleton. First, the kinematic model was used to perform a parameter sweep of the linkage dimensions to establish which combinations of dimensions could achieve the desired range of motion **(Supplementary Info. 4d**). Next, these combinations of dimensions were evaluated using the dynamic model to compute the motor torques and velocities that were required to achieve the desired pivot point (or mouse) velocities and accelerations (**Supplementary Info. 4e-f**). The optimal set of linkage dimensions were selected to produce low motor torque and velocity (**Supplementary Info. 4g**). Lastly, to return to ground truth, a time series of velocity and acceleration data obtained from marker-less tracking of freely behaving mice in an open field arena^9^ were input into the dynamic model using the optimal linkage dimensions. The resultant joint torques were parsed into gravitational, velocity, and acceleration components (**Supplementary Info. 4h**) and then these components were used to evaluate the linkage stiffness, mass, and material selection.

The admittance control law implemented in the force feedback controller instructs the exoskeleton to emulate user-programmable mass and damping values (the virtual admittance) that are orders of magnitude lower than its true values (the robot admittance). Because sensor sampling, computations, and feedback loops in the controller all introduce time delays, and motors have a finite torque limit, the accuracy of this emulation (the apparent admittance) degrades at higher frequencies, resulting in the mouse experiencing larger mass and damping values. To evaluate the limitations of the admittance control approach, we used a Laplace- space model of the exoskeleton and controller to identify the bandwidth of the system^23^ (**Supplementary Info. 5**). The bandwidth provides a measure of how quickly the exoskeleton can respond to force inputs from the mice before the apparent admittance degrades significantly. The bandwidth of the exoskeleton was estimated to be approximately 4 Hz, which was expected to produce a robot-rodent interface that was perceptible to mice but that would not hinder their volitional movement in navigation tasks.

### Rodent behavior-in-the-loop controller tuning to achieve natural locomotion

To achieve natural locomotion when mice are maneuvering the exoskeleton, we first evaluated the velocity and acceleration profiles of freely behaving mice (n = 3) in an open field arena from our previous study^9^. For short periods of motion, profiles generally followed arching trajectories with acceleration peaking around 50 to 75% of the peak velocity attained for the period of motion, before crossing the velocity axis and decelerating in a similar arching trajectory back to the origin (mouse stationary) (**Extended Data 1**, **Supplementary Video 1**). For sustained periods of motion, the trajectory arched towards a peak velocity and then followed smaller circular trajectories centered on the mean velocity. Ideally, the velocity and acceleration profiles of mice maneuvering the exoskeleton needed to match those observed in freely behaving mice, and these profiles needed to be achieved with mice exerting minimal forces. We tuned the admittance controller by starting with large mass (m) and damping (c) values to ensure stability and then incrementally decreased the m and c values until the velocity and acceleration profiles of mice maneuvering the exoskeleton were comparable to those observed in freely behaving mice (**Fig. 2a-b**). For all experiments, the pitch angle (20 degrees) and snout height (6 to 8 mm) for the exoskeleton were configured within the range of physiologically realistic values for mice at walking velocities, which we determined using marker-less tracking^24^ of freely behaving mice (n = 4) (**Extended Data 2**).

**Fig. 2:**
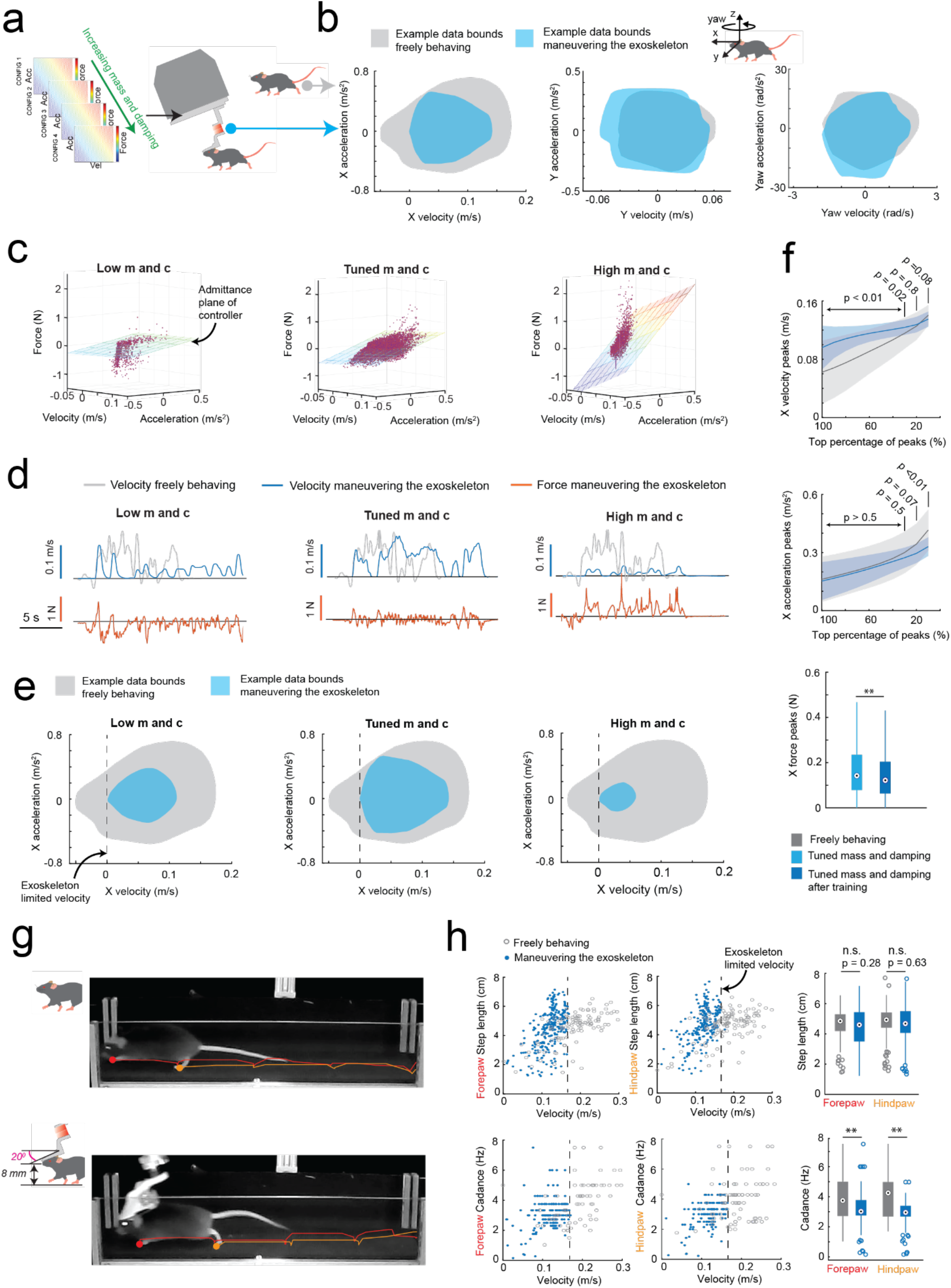
Rodent behavior-in-the-loop admittance controller tuning: (a) Methodology used for tuning the robot admittance controller to match locomotion characteristics of freely behaving mice. (b) Representative illustrations of velocity and acceleration data bounds in the mouse’s x (forwards), y (lateral) and yaw (rotation left and right) axes generated by a freely behaving mouse (grey) and by a mouse maneuvering the exoskeleton (blue). (c) Example raw data (purple points) of velocity, acceleration, and force in the mouse’s x axis with low (left), tuned (center), and high (right) mass and damping, overlaid with the admittance plane of the controller. (d) Representative velocity (blue) and force (orange) time series in the mouse’s x axis, generated by a mouse maneuvering the exoskeleton with low (left), tuned (center), and high (right) mass and damping, with velocity time series of a freely behaving mouse (grey) overlaid for comparison. (e) Representative velocity and acceleration data bounds in the mouse’s X axis generated by a freely behaving mouse (grey) and by a mouse maneuvering the exoskeleton (blue) with low (left), tuned (center), and high (right) mass and damping. (f) Comparison of the range of velocity (top) and acceleration (middle) peaks in the mouse’s x axis generated by freely behaving mice (grey) and mice maneuvering the exoskeleton with tuned mass and damping after training (dark blue); with p-values calculated using 1-way ANOVA. The distribution of force peaks (bottom) before (light blue) and after (dark blue) training. (g) Still images of the same mouse locomoting while freely behaving (top) and while maneuvering the exoskeleton (bottom), overlaid with the location history of the left forepaw and hind paw to qualitatively show similarity in step size. (h) Step lengths (top row) and cadence (bottom row) of the left forepaw and hind paw in freely behaving mice (grey) and mice maneuvering the exoskeleton (blue).

We tuned the mouse’s forwards direction (positive x axis) with a test cohort of mice (n = 4) maneuvering the exoskeleton around a linear oval track (**Extended Data 3**). In total, 14 combinations of m and c values were evaluated across 20 sessions. Large m and c values resulted in large amplitude, positive spiking forces, and discontinuous periods of motion at velocities and accelerations significantly less than those observed in freely behaving mice (**Fig. 2c-e** right). Decreasing the m and c values resulted in decreasing forces, increasing peak velocities and accelerations, and increasing periods of sustained motion. However, with the m and c values too low, motion became discontinuous again and the force became predominantly negative, indicating the exoskeleton was moving faster than the mouse intended (**Fig. 2c-e,** left). With optimally tuned m and c values, low amplitude, positive and negative forces were observed, and sustained periods of motion were achieved with velocities and accelerations comparable to those observed in freely behaving mice (**Fig. 2c-e**, middle, **Supplementary Video 1**). After training mice on the exoskeleton with optimally tuned m and c values, the mean peak forces decreased from 0.19 ± 0.06 N to 0.16 ± 0.03 N (**Fig. 2f** top). The range of velocities and accelerations became closer to those of freely behaving mice (**Extended Data 3e-f**), with the top 20% of velocity peaks of 13 ± 2 cm/s and 13 ± 1 cm/s (p = 0.07) and top 20% of acceleration peaks of 35 ± 10 cm/s^2^ and 30 ± 5 cm/s^2^ (p = 0.8) for freely behaving mice and trained mice maneuvering the exoskeleton, respectively (1-way ANOVA, p-values for full range of velocity and accelerations in **Fig. 2f**).

With the controller tuned, we evaluated gait dynamics to establish whether natural locomotion was being conserved in mice maneuvering the exoskeleton. An infrared camera captured the side-view of mice (n = 3) as they locomoted along a 27 cm long straight section in the linear oval track in both conditions (**Supplementary Video 2**). Representative still images of the same mouse locomoting when freely behaving and when maneuvering the exoskeleton, overlaid with traces indicating the left forepaw (red) and hind paw (orange) location history are shown in **Fig. 2g.** For all mice, the step length and cadence for both paws generally increased with their forwards velocity, in line with other studies^25^ (**Fig. 2h**). No significant difference in step length was observed between freely behaving mice and mice maneuvering the exoskeleton, with mean forepaw and hind paw step lengths of 4.6 ± 1.1 and 4.7 ± 1.1 cm in freely behaving mice, and of 4.5 ± 1.3 and 4.7 ± 1.2 cm in mice maneuvering the exoskeleton (**Fig. 2h**; 1-way ANOVA; forepaw: p = 0.28; hind paw: p = 0.63). However, the cadence did vary significantly, with mean forepaw and hind paw cadence of 3.9 ± 1.5 and 4.1 ± 1.5 Hz in freely behaving mice, and of 3.1 ± 1.0 and 3.0 ± 0.8 Hz in mice maneuvering the exoskeleton (**Fig. 2h**; 1-way ANOVA; forepaw: p = 0.28; hind paw: p = 0.63). We suspect that the observed difference in cadence is partially caused by an upper velocity limit of 16 cm/s that was implemented on the exoskeleton controller, and partially caused by the bandwidth of the exoskeleton, the mass and damping values of the admittance controller, and the constant pitch and snout height.

### Mice maneuvering the exoskeleton make turns and navigate 2D arenas

Having established that mice could locomote naturally in their forwards direction while maneuvering the exoskeleton, we next provided mice with control of their lateral (y axis) and turning (yaw axis) directions so that they could make turns. In an 8-maze arena, mice had control of their x, y, and yaw axes in a central zone and had motion in the two goal arms confined to their x axis along a circular trajectory (**Fig. 3a-b, Extended Data 5**). We first used this arena to tune the y and yaw axes using the same methodology as the x axis, with a total of 7 combinations of m and c values in the y axis and 6 combinations in the yaw axis evaluated across 15 sessions in a test cohort of mice (n = 2) (**Extended Data 4**). Once the y and yaw axes were tuned, we took a new cohort of mice (n = 5) with no experience on the exoskeleton and evaluated their turning proficiency across several training sessions in the 8-maze arena.

**Fig. 3:**
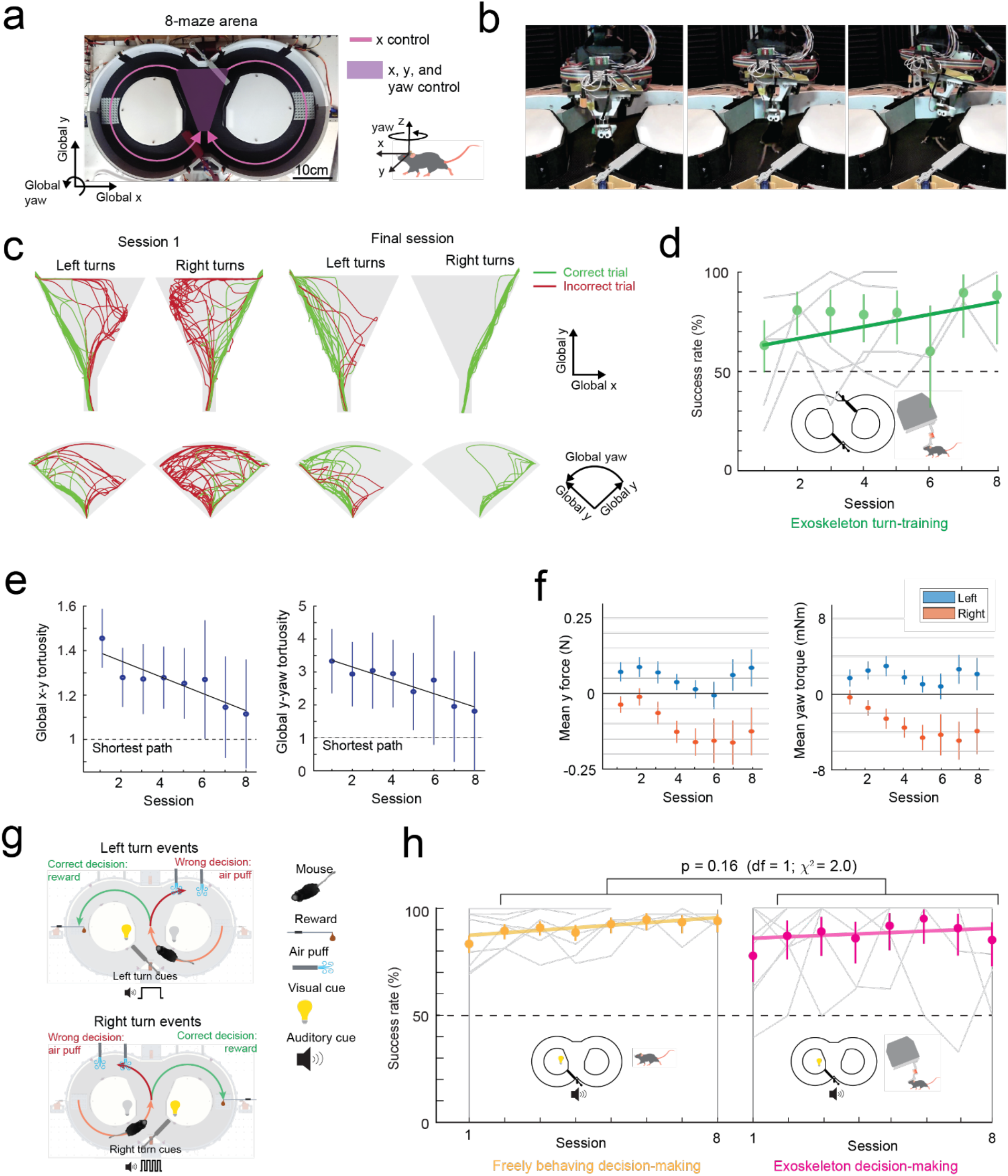
Mice maneuvering the cranial exoskeleton can perform a navigational decision- making task. (a) Photograph of the 8-maze arena overlaid with the exoskeleton control implementation, Mice controlled their x axis motion in the arms of the maze (pink arrows) and had control over their x, y, and yaw axes motion in the turning zone (purple region). (b) Series of images showing a mouse turning right in the turning zone. (c) Paths traversed by mice through the turning zone (n = 5 mice) on their first (m = 57 turns) and final (m = 55 turns) sessions. Green – correct turns, Red – incorrect turns. (d) Performance of mice (n = 5) in executing turns in the turning zone while maneuvering the exoskeleton. (e) Mean and 95% confidence intervals of the global x-y (left) and global y-yaw (right) path tortuosity through the turning zone for all mice (n = 5) across 8 sessions. (f) Mean and 95% confidence intervals of the y (left) and yaw (right) forces during left (blue) and right (orange) turns for all mice (n = 5) across 8 sessions. (g) Schematic of the navigational decision-making task showing events for left (top) and right (bottom) turns. (h) Performance of mice (n = 8) in the navigational decision-making task while freely behaving (left, yellow) and while maneuvering the exoskeleton (right, magenta).

We mapped the paths the mice took through the turning zone in the global x-y axes and the global y-yaw axes and then categorized the turn as an incorrect trial if the global x-y path deviated by more than 1 cm from the midline in the contralateral direction to the turn (**Fig. 3c**). Using this scoring method, all mice achieved greater than 80% turning proficiency within 8 sessions, with 2 mice achieving the same proficiency after 3 sessions (**Fig. 3d**). The corresponding tortuosity of the tracked paths within the turn zone in global x-y and global y-yaw paths both showed a decreasing trend across sessions as mice took more direct paths between the entry and exit points of the turning zone (**Fig. 3e**). The magnitude of the difference between left and right turning forces increased across sessions (**Fig. 3f**, **Extended Data 6**). These trends in the tortuosity and forces suggest that once mice were familiar with the robot-rodent interface they could consistently and proficiently perform turning while maneuvering the exoskeleton.

### Mice maneuvering the exoskeleton can perform a navigational decision-making task

We next asked whether mice that can turn proficiently in the 8-maze arena could also make navigational turning decisions. In these experiments, the door at the exit of the turning zone (training door) was removed so that mice were free to select either goal arm, and visual and auditory cues were introduced that were unique to the correct turn decision (left turn cues: light on left side of the turning zone and 1 long auditory tone 5 kHz, 4 s duration; right turn cue: light on the right side of the turning zone and 4 short auditory tones 5 kHz, 0.75 s ON / 0.25 s OFF) (**Fig. 3g**, **Supplementary Info. 7**). A cohort of mice (n = 8) were first trained while freely behaving to alternate between left and right turns, receiving a reward for each correct decision and an air puff for each wrong decision. Once trained, mice consistently achieved a mean success rate of 88 to 95% (1^st^ session excluded), with 4 mice achieving 100% success rate in at least 1 session for a combined total of 10 of the 50 sessions (**Fig. 3h**). These mice were then docked to the exoskeleton in the 8-maze arena, where they performed the same navigational task with a mean success rate of 85 to 95% (1^st^ session excluded), and with all 8 mice achieving 100% success rate in at least 1 session for a combined total of 37 of the 60 sessions, (**Fig. 3h**, **Extended Data 7, Supplementary Video 3**). There was no statistically significant difference in task performance between the exoskeleton maneuvering and freely behaving scenarios (Kruskal-Wallis test, p = 0.16, 1^st^ session excluded from both groups) (**Fig. 3h, Extended Data 7**), however the variability in performance was larger in mice maneuvering the exoskeleton than in freely behaving mice (Clopper-Pearson binomial: ±2% freely behaving, ±10% maneuvering exoskeleton). These results demonstrate that mice readily learn to navigate 2D physical spaces while maneuvering the exoskeleton and can perform learned cognitive tasks, such as sensory cue-guided decision-making, at the same performance level as when they are freely behaving.

### Ultra-widefield cellular resolution imaging of the cortex during 2D navigation

In the first of two headstage designs that leverage the payload carrying capabilities of the exoskeleton, we incorporated a simple, custom built macroscope for widefield imaging (**Fig. 4a**). The macroscope used a 1x zoom macro-lens coupled to a 5-megapixel, monochrome camera to image an 8.6 x 6.6 mm field of view at a design resolution of 3.45 x 3.45 µm (pixel size) (**Supplementary Info. 2**). In bench-top imaging of resolution test-targets, line widths in the range of 3.5 to 5.5 µm could be resolved across the field of view (**Fig. 4b**). Double transgenic mice^26^ (Cux2-Cre-ERT2^27^ X Ai163^28^, n = 4 mice) sparsely expressing the Ca^2+^ indicator GCaMP6f in layers II-III excitatory neurons were implanted with a planar glass window making 35 mm^2^ area of the dorsal cortex optically accessible for imaging (**Fig. 4a, c**). High resolution mesoscale imaging data was acquired in each session that mice navigated the 8-maze arena.

**Fig. 4:**
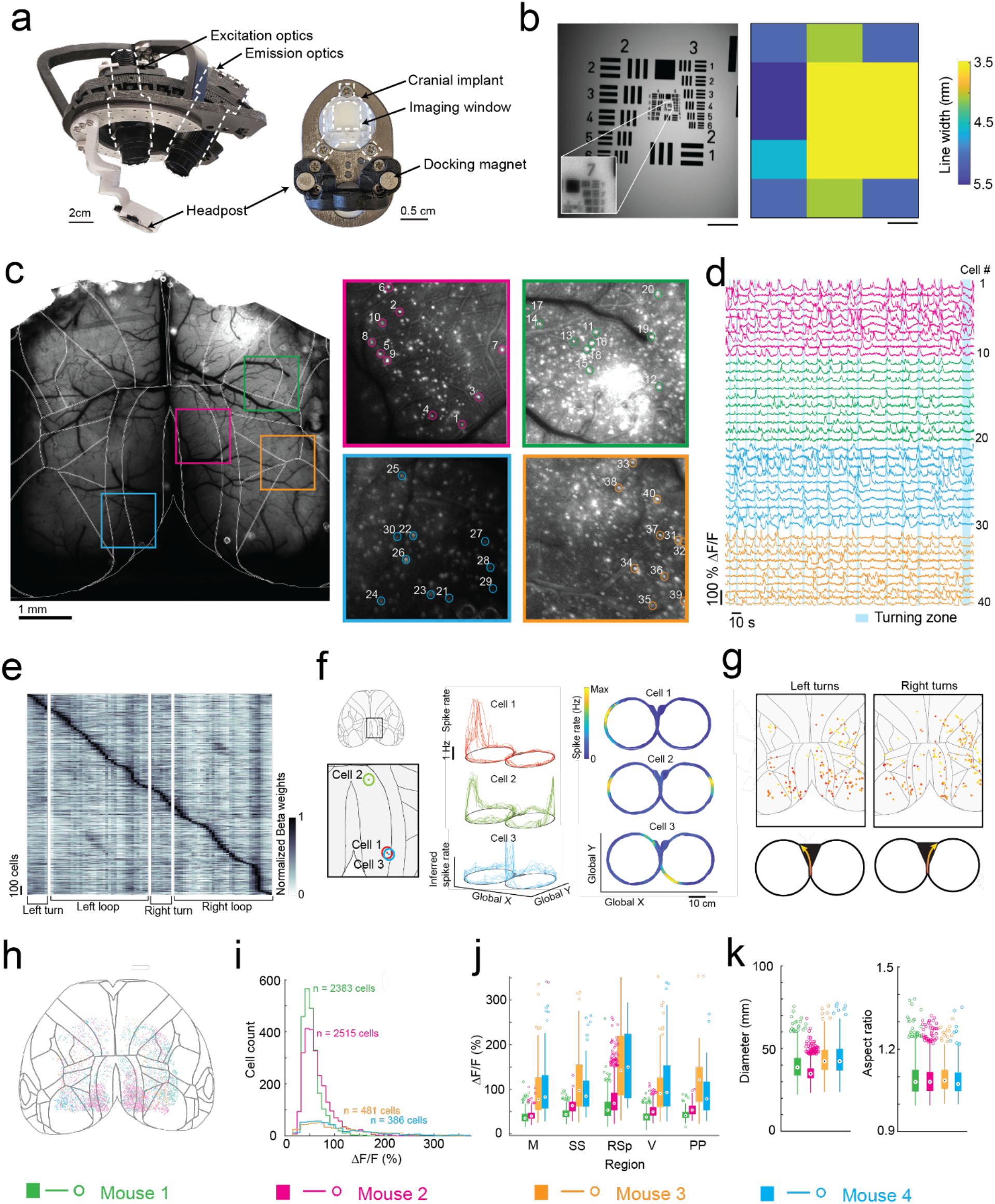
Mesoscopic cellular-resolution cortical imaging in mice navigating an 8-maze arena. (a) Photographs of the headstage (left) and headpost (right) for mesoscale cellular-resolution imaging. (b) Image of a resolution test target (left) acquired using the headstage with (inset) digital zoom showing the resolution limit at group 7-2, and (right) variability in resolution across the field of view. Scale bars are 1 mm. (c) Raw image of the dorsal cortex with brain-region boundaries overlaid, and maximum intensity images of 4 regions of interest (ROI; 1x1 mm) with 10 cells in each ROI numbered 1 through 40. (d) Raw fluorescence traces (ΔF/F) of cells 1 through 40 and bands (light blue) showing when the mouse was in the turning zone. (e) Kernel matrix of normalized Beta weights obtained using linear regression on the inferred spike rate of each cell (n = 2383 cells) and the location of the mouse in the 8-maze arena (m = 82 location bins), with rows in the Kernel sorted by onset of the maximum beta weight. (f) Inferred spike rates of 3 cells in the retrosplenial cortex that each fired at specific locations in the 8-maze arena (cell 1, left arm; cell 2, left and right arms; cell 3, before and after left turns in the turning zone). (g) Locations of all cells with normalized Beta weights of 1 occurring during left or right turns, with color scale indicating progression through the turning zone (starting in red and ending in yellow). (h) Anatomical locations of all cells imaged across all mice (n = 4) overlaid on the Allen brain atlas reference map. (i) Histogram showing the cell fluorescence (ΔF/F) distribution of all cells for each mouse. (j) Break-down by region of the cell fluorescence (ΔF/F) distribution for each mouse. (k) Distribution of soma diameters (left) and aspect ratios (right) for each mouse.

The mesoscale imaging headstage allowed us to reliably record Ca^2+^ activities of 1000s of neurons from several regions distributed across the cortex (**Fig. 4c, Extended Data 8a-e, Supplementary Video 4**). Example calcium fluorescence (ΔF/F) traces from 40 neurons recorded from 4 disparate brain regions in one mouse (mouse 1, session 9) completing 16 trials in the 8-maze arena while maneuvering the exoskeleton are shown in **Fig. 4d**. After applying exclusion criteria on the region of interest (ROI) morphology and signal amplitude (see Methods), we identified 2383 cells that could be simultaneously recorded in this mouse.

To evaluate spatial encoding of the 8-maze arena, we performed a linear regression analysis on the inferred spiking activity and the mouse’s location within the arena. This generated a kernel matrix of Beta weights relating each cell’s firing rate to the location. Sorting the normalized Beta weights of each cell by the onset location of their maximum produced a clear diagonal trend indicating many of the cells preferentially fired at certain locations, as well as vertical bands in some locations indicating increased global activity (**Fig. 4e, Extended Data 8f**). Examples of this preferential spatial firing are shown in **Figure 4f** for 3 cells in the retrosplenial cortex, where cell 1 fired at a tactile feature (Lego^TM^ floor) in the left arm of the 8-maze (**Fig. 4f** top), cell 2 fired in both left and right arms over the same tactile feature (**Fig. 4f** middle), and cell 3 fired before and after left hand turns through the turning zone (**Fig. 4f** bottom). Selecting the cells with maximum Beta weights through the turning zone revealed distinct populations of neurons distributed across the whole cortex that preferentially fired as the mouse made left and right turns through the turning zone (**Fig. 4g**).

In total, we recorded 5,765 cells distributed across the cortices of 4 mice (**Fig. 4h-j, Extended Data 8**). Cell somas were on average 11 ± 2 pixels in diameter (38 ± 7 µm) and had an aspect ratio of 1.1 ± 0.1 (**Fig. 4k**). Importantly, moving the imaging hardware along with mice did not cause significant motion artifacts. Lateral motion artifacts could be corrected for using the standard rigid body motion correction algorithm applied within the cell sorting package Suite2p^29^ with maxima of 18 pixels (62 µm) and 15 pixels (52 µm) motion correction applied in the anterior-posterior and medial-lateral directions respectively. Variability in expression patterns were observed across mice, with increased sparsity in expression corresponding to much brighter GCaMP expression in individual cells (**Fig. 4h-i**). In two mice, 2383 and 2515 cells were identified with smaller average peak ΔF/F of 45 ± 18% and 60 ± 29% respectively, whereas in the other two mice 386 to 481 cells were identified with larger average peak ΔF/F of 112 ± 72% and 108 ± 68%, with cells across all 4 mice identified using the same thresholds^29^ (see Methods). This variation in expression patterns is likely due to differential expression of GCaMP6f unlocked via tamoxifen treatment. These experiments reveal the information-rich, large-scale neural activity datasets that can be reliably and uniquely acquired with the cranial exoskeleton as mice explore physical spaces.

### Multi-day, multi-site brain wide electrophysiology recordings during navigational decision-making task

We engineered a second headstage for the exoskeleton to perform multi-site electrophysiology (**Fig. 5a**). The headstage incorporated stacks of 3-degrees-of-freedom (DOF) micro- manipulators for simultaneously aligning and inserting up to 4 neural probes, and a miniature behavioral camera with infrared-vision to acquire video of the mouse’s head and forepaws (**Supplementary Info. 3**). Mice (n = 4) were implanted with 3D printed, polymer, cranial implants, modified from our previous work^8,9^, which contained 6 probe-entry ports providing physical access for neural probes to the brain (**Fig. 5b**, **Supplementary Info. 8**). These mice were trained on the navigational decision-making task (sensory-cue guided alternating-choice) in the 8-maze arena until they reached a proficiency of >85% for at least 2 consecutive sessions (**Fig. 3**) before acquiring electrophysiology recordings in the task. Immediately prior to recordings, mice were docked to the exoskeleton in a staging area where they could run on a disk-treadmill as probes were aligned and inserted into the brain (**Supplementary Video 5**).

**Fig. 5:**
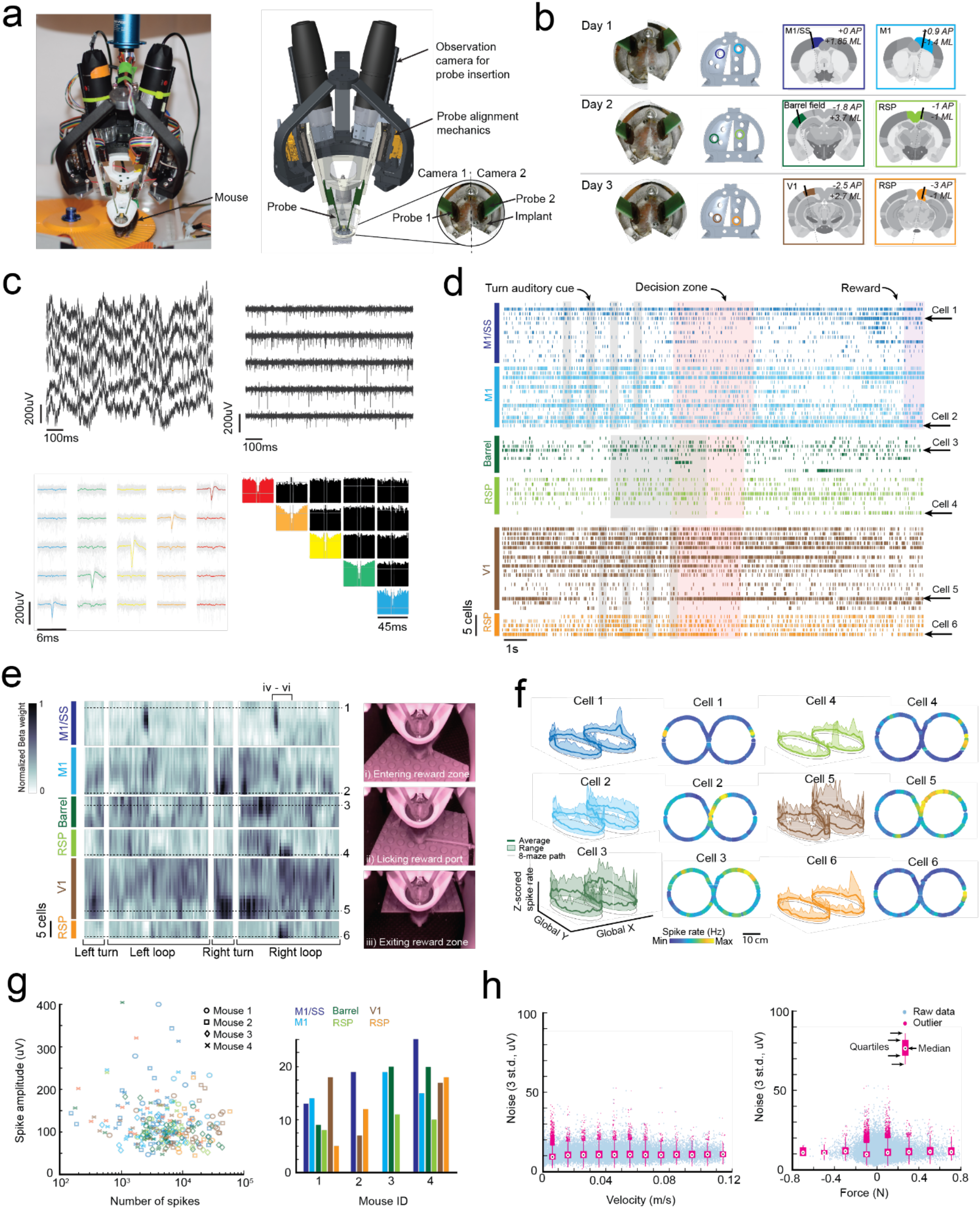
Multi-site, multi-day electrophysiology recordings across the brain in mice performing a navigational decision-making task. (a) Photograph (left) and CAD rendering (right) of the headstage for electrophysiology with observation cameras mounted to monitor probe alignment and insertion. (b) Images of inserted probes (left), the corresponding probe entry ports on the chronic implant (center-left), and the target brain regions across the 3 recording days (center-right and right); AP, Anteroposterior; ML, Mediolateral. (c) Example data showing raw traces recorded from 5 neighboring electrodes on a neural probe before (top-left) and after (top-right) noise removal, and the mean waveforms (bottom-left) and correlograms (bottom-right) of 5 cells identified on these electrodes. (d) Spike raster of all cells recorded from 1 mouse across 3 days, with auditory cues (grey), turning zone (red) and reward port (purple) events overlaid. (e) Kernel matrix of normalized Beta weights obtained using linear regression on the z-scored spike rate of each cell (n = 67 cells) and the location of the mouse in the 8-maze arena (m = 196 location bins). (f) Average z-scored spike rates of 1 cell from each probe insertion site showing activity patterns across the 8-maze arena (cell 1, left and right rewards; cell 2, right turns through turning zone; cell 3, left and right arms; cell 4, right arm after reward; cell 5, right turns through turning zone; cell 6, right arm after reward). (g) Spike amplitudes of all cells with colors indicating insertion site and shapes indicating the mouse. (h) Number of spikes and spike amplitude for each cell, with same color and shape identifiers as (g). (i) Distribution of noise (3 standard deviations) with respect to mouse velocity (left) and force (right).

Following probe insertion, mice began the task by entering the 8-maze arena along a linear trajectory into the start of the turning zone (**Extended Data 5b**).

A total of 18 recordings were acquired across 10 recording sessions in the 4 mice performing the decision-making task, with example data and analysis from 3 sessions in 1 mouse (mouse 1) shown in **Fig. 5c-f**. Any motion artifacts that occurred, either due to relative motion of the kinematic clamp with respect to the fixed probes in the headstage or relative motion of the brain with respect to the cranial implant, were corrected using the standard motion correction algorithm applied within the spike sorting package Kilosort 2.0^30^. Bandpass filtering and common-mode subtraction reduced the 3 standard deviation noise on the raw voltage traces to an average of 10 ± 5 µV (**Fig. 5h**) indicating that any environmental noise introduced by the robotic elements of the exoskeleton can be removed using standard noise reduction techniques. Spiking signals were clearly visible in raw traces after noise reduction (**Fig. 5c**), and manual curation of the spiking signal waveforms^31^ identified multiple well-isolated cells on each probe (**Fig. 5c-d, Extended Data 9a-b**). The number of cells identified on each probe varied from 5 to 25 due to the depth of probe insertion and the electrode spacing (50 or 100 µm). Regular-, fast-, triphasic-, compound-, and positive-spiking waveforms were all present across the recordings, with amplitudes ranging from around 50 and 400 µV (**Extended Data 9b-c**).

In addition to the decision-making, turning, and reward port-licking behaviors that were expected in the navigational decision-making task, mice also exhibited natural behaviors during recording sessions such as grooming their snout and gripping the tactile features on the Lego^TM^ floor (**Fig. 1c-d**, **Fig. 5e**, **Supplementary Video 5**). A linear regression analysis, similar to that performed on the single-cell imaging data (**Fig. 4e**), was performed on the 2 mice with full sets of recordings (6 probes across 3 sessions each) to evaluate spatial encoding of the 8-maze arena in the cell spike rates (**Fig. 5e**, **Extended Data 9d-e**). The resulting kernel matrices were normalized for each cell and were arranged in order of descending cortical depth for each recording site (top row of the kernel closest to cortical surface). Preferential spiking was observed in the kernel (**Fig. 5e**) and in cell firing patterns (**Fig. 5f**) for mouse 1 over the tactile floor features on both arms (cell 1, M1/SS), during right hand turns through the turning zone (cells 2 and 5, M1 and V1), at the start of the left and right goal arms (cell 3, Barrel), and after the reward port in the right arm (cells 4 and 6, RSp). We recorded from a total of 261 cells during the decision-making task, with the number of spike signals in each cell ranging from 150 to 60,000 spikes (average of 11,900 spikes) recorded across sessions lasting between 5 and 33 minutes (16 ± 9 minutes) (**Fig. 5g**). Importantly, we saw no correlation between the signal noise amplitude and either increasing mouse velocity or increasing mouse force, indicating that moving the recording hardware along with mice did not cause significant motion artifacts (**Fig. 5g**). These experiments demonstrate that highly sensitive electrophysiology recordings can be reliably acquired across multiple recording sessions using the cranial exoskeleton as mice perform behavioral tasks.

## DISCUSSION

In summary, we report here a cranial exoskeleton that allows mice to navigate 2D physical environments and perform navigational decision-making tasks while large payloads (1 to 2 kg) of equipment acquire brain-wide neural recordings. To accomplish this, we realized a robot-rodent control interface that enables mice to locomote at physiologically realistic velocities and accelerations in their x, y, and yaw axes and conserves natural gait along straight trajectories. We also realized two neural recording headstages capable of performing brain-wide recordings as mice navigate physical spaces; one headstage for wide-field, cellular resolution imaging of 1000s of neurons distributed across the cortex, and a second headstage for multi-site, multi-day electrophysiology using silicon probes. The hardware and techniques implemented on our wide- field imaging and multi-site electrophysiology headstages are already used in modern neuroscience^11, 32^ but until now have been limited to studies in head-fixed subjects, often incorporating virtual reality to investigate navigational elements of behavior. The exoskeleton allows many of these head-fixed studies to be expanded to physical environments.

Physical environments produce a more natural, multi-sensory experience for mice by incorporating motion-vestibular input coupling, natural eye-head movement coupling, whisking of physical boundaries, and natural spatial representations of sensory information (for example varying sound, light, and odor intensities with proximity to source, or shadowing effects of objects). Consequently, the neural representations in actual physical environments should be more natural and multi-sensory, providing avenues for new exploration into how brain-wide neural activity mediates ethologically relevant and complex behaviors. Importantly, the ability of the exoskeleton to control aspects of the experiment design provides researchers with the capabilities to control elements of the mouse’s experience which can otherwise be difficult in freely behaving experiment designs. This retention of precise experimental control should provide researchers with the tools to unravel the neural representations of higher cognitive functions such as multi-sensory integration, decision-making, and 2D or 3D navigation.

In the current study, mice were restricted to either 1 (x) or 3 (x, y, yaw) DOF of control in the behavioral tasks as we built an understanding of robot-rodent interaction dynamics. Moreover, the mass and damping values that the mouse experienced in these axes were not imperceptible due to the relatively low bandwidth (4 Hz) of the robot-rodent interface. While these limitations did not affect decision-making performance in the navigational task that we studied, they were expected to have altered some behaviors, such as gait during turning, and possibly even suppressed other behaviors altogether, such as darting and rapid head turning^17, 33, 34^ which would require a faster response from the exoskeleton controller. Increasing the exoskeleton to full 6 DOF control and increasing the bandwidth to achieve a more imperceptible interface are expected to permit more natural representations of some of the behaviors that we observed, such as grooming and turning, as well as a broader range of behaviors overall.

The admittance controllers we used in this study were linear, which provided a predictable robot-rodent interface for mice to familiarize with. Non-linear control approaches, customization of control parameters for each mouse, and gradually varying the control parameters as mice are trained, are all avenues for exploration that may improve the overall robot-rodent interface.

Sophisticated control laws such as model-predictive control have been successfully implemented on robotic exoskeletons for humans^35, 36^ and may prove beneficial in rodent exoskeletons provided suitable models of mouse behavioral kinematics and dynamics can be defined. Within any control model, including linear admittance control, the location of the virtual pivot point about which the headstage is manipulated may not be constant across behaviors.

Moreover, while the forces are measured through the chronically implanted headpost, they are often generated through the mouse’s legs and body and so interpretation of the mouse’s intentions will also be influenced by the mouse’s stance, body orientation, and pose. Therefore, realizing a more complete range of natural behaviors on mice maneuvering an exoskeleton may require monitoring of several body points in addition to the cranial forces, an approach known as sensor-fusion.

The electrophysiology headstage weighed 1 kg and the imaging headstage weighed 1.5 kg: both far below the 3kg design load carrying capability of the exoskeleton. An array of high-speed, multiple perspective, behavioral cameras mounted to the headstage, and suitably positioned infrared mirrors, would provide comprehensive tracking of body-wide behavioral metrics. Future headstage designs for simultaneously inserting up to 12 or more CMOS recording probes can be incorporated that will allow simultaneous recordings from 10s of 1000s of neurons at a time. Similarly, other modalities, such as multi-photon mesoscale imaging systems^5, 6, 37^, combined electrophysiology and optogenetics^38^, and patterned stimulation^39^, that would be impossible to implement in a miniaturized configuration^40^ can be implemented on an exoskeleton to allow for comprehensive mapping or modulation of brain-wide neural activity.

## METHODS

### Exoskeleton construction

#### Exoskeleton Design

Within the kinematic model, we calculated the delta robot kinematics using the geometric method^41^ with joint angle limits set to +45 degrees and -90 degrees, and goniometer kinematics using standard coordinate transform techniques for serial robots^42^ (**Supplementary Info. 4b**). The Jacobian elements for the delta robot were calculated using the vector method^43^, and the elements for the goniometer and virtual pivot point were calculated using Euler matrix factorization^42^. Within the dynamic model, the delta robot dynamics was based on a model by Zhang et al^43^ with an additional force term acting on the delta robot moving platform that described dynamic forces generated by the goniometer. The dynamics of the goniometer was modelled using the Recursive Newton-Euler Algorithm^44^ with the velocity and acceleration of the delta robot moving platform assigned to the goniometer reference frame to couple it to the delta robot dynamic model (**Supplementary Info. 4e**).

#### Exoskeleton construction

The key dimensions of the constructed delta robot were a fixed platform radius (*R*_*f*_) of 0.19 m; moving platform radius (*R*_*m*_) of 0.1 m; proximal link arm length (*L*_*p*_) of 0.34 m; and distal link arm length (*L*_$_) of 0.656 m (**Supplementary Info. 1a-c and 4b**). The common rotation points of the three goniometer axes (*a*_1_) was offset 0.1 m below the center of the delta robot moving platform, and the virtual pivot point extended a further 0.25 m (*a*_4_) from this rotation point. All actuated joints in the exoskeleton were driven by backlash-free harmonic gear-drive servo-motors. The delta robot motors (FHA-17C-50-US250-E-SP, Harmonic Drive LLC) were capable of 30 Nm maximum torque and 576 deg/s maximum velocity, whereas the goniometer motors (FHA-8C- 100-US200-E, Harmonic Drive LLC) were capable of 4.8 Nm maximum torque and 360 deg/s maximum velocity. The delta robot link arms were constructed from carbon fiber to produce a high stiffness to weight ratio. The link arm joints at either end of the distal link arms were custom designed and contained two orthogonal revolute joints with intersecting axes of rotation to simplify kinematics and construction. The link arm joints were constructed from 3D printed clevis forks (Rigid10K, Formlabs) that were coupled to rotary shafts via a series of shaft couplings and ball bearings and contained a single-wave washer and thrust bearing to ensure high stiffness and low friction joint operation. The delta robot moving platform and goniometer motor brackets were fabricated from lightweight aluminum (Al 6061), whereas the delta robot fixed platform and delta robot motor brackets were fabricated from carbon steel plate (ASTM A36). The whole exoskeleton was mounted to a structural frame built using aluminum extrusion (15 series, Grainger) which was housed in a sound-proof enclosure.

For complete CAD design files and parts list of the exoskeleton, frame, and sound-proof enclosure, see the GitHub repository: https://github.com/bsbrl/exoskeleton

#### Control hardware

The delta robot and goniometer 3-phase motor inputs and encoder sensor outputs were connected to motor servo-drivers (DDP-090-36 for delta robot; DDP-090-09 for goniometer; Harmonic Drive LLC) which performed high speed current and velocity proportional-integral-derivative (PID) control. Motor and sensor cables for the goniometer were routed along the delta robot arms to the servo-drivers using high-flex cabling intended for repeated bending and torsion (T1376-5-ND, Digikey; 839-30-01159-30-ND, Digikey). The servo drivers were powered using a 24V DC power supply (B07WLKYNSH, Yi Mei Da via Amazon) capable of supplying 40A of current. The servo-drivers interfaced with a CompactRIO controller (NI-9038, National Instruments) which performed all kinematics, admittance control, and safety functions within its onboard software (LabVIEW 2019, National Instruments). The kinematics and admittance control calculations on the CompactRIO convert joint position and force sensor signals into target joint velocity signals (**Supplementary Info. 5 and 6**). The CompactRIO controller received the joint positions in the form of emulated differential quadrature encoder signals output from the servo-drivers. These differential signals were passed through dual differential line receivers (SN75157P, Texas Instruments) to convert them to single ended (ground referenced) signals before being input into high-speed encoder counters on the CompactRIO (NI-9401, National Instruments). Non-contact position referencing of the delta robot was achieved using magnets (469-1005-ND, Digikey) placed on the delta robot arms and hall effect switches (480-2006-ND, Digikey) placed on the delta robot fixed platform and were measured using the CompactRIO analogue input module (NI-9205, National Instruments). A CompactRIO analogue input module (NI-9205, National Instruments) measured 12 analog signals (6 differential pairs) output from the force sensor pre-amplifier electronics (9105-IFPS-1, ATI Industrial Automation) via shielded cable (9105-C-PS-U-2, ATI Industrial Automation). The target joint velocity signals were output from the CompactRIO (NI-9401, National Instruments), in the form of 10 kHz PWM signals, with 50% duty cycle equating to a target velocity of 0 encoder counts/second, which were input to the servo-drivers. Synchronization between the controllers and data acquisition systems was implemented using 3.3V digital pulses output from the CompactRIO controller (NI-9403, National Instruments) that were voltage buffered and amplified using a MOSFET (IRL520-ND, Digikey) connected to a 5V bench-top power supply. In behavioral experiments, light and sound cues, solenoid activated air puff and reward dispensing, and servo-actuated doors were also operated by the CompactRIO controller (NI- 9401, NI-9403, National Instruments), **Supplementary Info. 7**.

### Headstage construction

#### Headstage overall

The headstage was coupled to the goniometer through a custom made through-rod machined from Al 6061. A slip ring commutator with aperture was fitted around the outside of the through- rod (EM022-24GG, Senring). Three struts, 3D printed from fiber reinforced nylon (Markforged Mark 2), joined the headstage coupling to the mounting platform, which was waterjet cut from 5 mm thick Al 6061 plate and contained a hole pattern tapped M3 for mounting experiment hardware. A 3D printed strut (Rigid10k, Formlabs) extended down from the underside of the mounting platform to a 6-axis force-sensor (9105-TW-NANO17-E-2.5 with SI-25-0.25 calibration, ATI Industrial Automation). A 3D printed kinematic clamp (Rigid10k, Formlabs) was attached to the opposite side of the force sensor and contained two magnets (2455-ALC4010- ND, Digikey) to facilitate quick docking of mice, and two M5 ball-tip setscrews to lock mice in place via their chronically implanted headpost. The headpost contained a 3D printed docking mechanism (Black v4, Formlabs) with 2 magnets and 2 kinematic coupling features that mated to the kinematic clamp. The docking mechanism was fixed using four 0-80 screws to a 1mm thick titanium plate (ASTM B265) that kept the structure rigid, and the titanium plate was fixed using three 0-80 screws to the cranial implant that was chronically implanted on each mouse. (**Supplementary Info. 1**).

For complete CAD design files of the headstage and headpost see the GitHub repository: https://github.com/bsbrl/exoskeleton

#### Imaging headstage

The headstage for mesoscale imaging contained a 3D printed bracket (Stratasys Objet500) with an adjustable mounting system to support the excitation and emission optics (**Supplementary Info. 2**). The excitation optics component-stack contained a 470nm mounted LED (M470L5, Thorlabs) coupled to an adjustable collimator (SM1U25-A, Thorlabs) via a cage plate (CP02/M, Thorlabs) that was also used to mount the component stack to the headstage. A bandpass filter (MF469-35, Thorlabs) with 469 nm center wavelength and 39nm bandwidth was attached to the optical output end of the adjustable collimator using a filter mount (#65-800, Edmund Optics) and 2 thread adaptors (SM1A38 & SM1A24, Thorlabs). The LED was operated using a bench- top LED driver (LEDD1B,Thorlabs) that was triggered using a 5V digital TTL signal generated by the exoskeleton control hardware. The emission optics component stack contained a monochrome, 2/3” format, 5.07 megapixel camera (Allied Vision Alvium 1800 U-508m: #17-087, Edmund Optics) attached to a macro lens with manually adjustable zoom (1x to 3.3x), focus, and f/# (#56-524, Edmund Optics). A bandpass filter (MF525-39, Thorlabs) with 525 nm center wavelength and 39nm bandwidth was attached to the optical input end of the macro lens using the same type of filter mount and thread adaptors as the emission filter.

#### Ephys headstage

The headstage for multi-site electrophysiology contained a 3D printed bracket (fiber reinforced nylon, Markforged Mark 2) that could support docking of up to 4 neural probes and accompanying probe alignment mechanics (**Supplementary Info. 3**). In the current study, only 2 probe docking sites were used at a time. The probe alignment mechanics contained a linear axis (M3-LS-3.4-15, New Scale Technologies) with 15 mm travel for lowering the probes into the brain, and two angular axes (9061-PY-M, Newport) with ±2 degrees travel range for controlling the lateral position of probe tip. The neural probes (A1x32-Edge-5mm-50-177-A32 or A1x32- Edge-5mm-100-177-A32, NeuroNexus) and head stage electronics (#C3314 RHD 32ch, INTAN) were attached to the probe alignment mechanics using an adaptor (Adpt-A32-OM32, NeuroNexus) mounted to a 3D printed bracket (Rigid10k, Formlabs). Ground connection to the mouse was made by connecting a wire from the head stage electronics ground contact and a wire from the mouse’s ground skull-screw to a screw terminal block located on the headstage near to the mouse.

#### Behavior imaging

Behavioral imaging was acquired using cameras placed around the behavioral environments and on the headstage. The cameras placed around the behavioral environments were either webcams (C920S Pro HD 1080p, Logitech) or night vision cameras with infrared LED arrays (ELP-USBFHD01M-DL36, SVPRO) and were operated using a host PC. The camera on the headstage was a miniature night vision camera with 2 infrared LED lamps (1778-1218-ND, Digikey) that was operated using a Raspberry Pi (2648-SC0510-ND, Digikey) (**Supplementary Info. 3**). The Raspberry Pi was powered using a 5V USB power bank mounted on the headstage. The camera on the headstage was directed at the mouse’s head and captured an approximately 100 x 80 mm field of view that encompassed the headpost, snout, whiskers, and front paws of the mouse, as well as up to 2 infrared mirrors (1601-G380227033-ND, Digikey) that were angled to capture the mouse’s eyes. The infrared mirrors were used in several development experiments but were not used in the 8maze behavioral experiments.

### Controller

#### Control diagram

The controller contained 3 PID control-loops in series: a position loop that received target and actual operational-space positions and output a target operational-space velocity; a velocity loop that received target and actual joint velocities and output a target motor current; and a current loop that received target and actual motor currents and output an actual current to the motors (**Supplementary Info. 5**). The current and velocity loops were located on the motor servo- drivers, whereas the position loop was located on the CompactRIO controller. The currents supplied to the motors generated torques in the joints, that produced changes in the joint velocities, which were fed into the velocity loop, and changes in the actual operational-space position. The forces applied by the mouse to move the exoskeleton were measured by the force sensor and fed into an admittance controller which, together with the current operation-space velocity, calculated a target operational-space acceleration. This acceleration was then double integrated with respect to time to find a target position that was fed into the position loop (**Supplementary Info. 5 and 6**).

#### Tuning

The PID control-loops were tuned one at a time starting with the current loop and then moving backwards through the control diagram. The current loop was tuned using an automated tuning feature in the software provided by the supplier (HDM Software v71B24, Harmonic Drive LLC). The velocity loop was manually tuned within the same software by optimizing the step and settle response to small amplitude 1 Hz square wave inputs and the tracking error of 5 Hz sine wave inputs. Before tuning the position loop, the open-loop bandwidth in operation-space was characterized by commanding sinusoidal velocity profiles at frequencies from 0.1 to 10 Hz. The bandwidth was determined to be 4.5 Hz using a Bode plot (phase and magnitude) of the error between the commanded operational-space velocity and the resultant operational-space position. The position loop was then manually tuned in each axis by optimizing the step and settle response to small amplitude 1 Hz square wave. The z axis was tuned more aggressively than the x and y axes because this axis was fixed in the current study, and so the errors were expected to be comparatively small.

#### Bandwidth

A state space model of the delta robot and controller was used to identify and improve bandwidth constraints^23^. An example of one improvement that was implemented is shown in (**Supplementary Info. 5**) where optimizing code on the CompactRIO controller reduced the cycle time from 50 ms to 10 ms, which resulted in a 2- to 4-fold improvement in bandwidth. Other improvements that were implemented include a reduced temporal delay of the force sensor signal (zero order hold component in the state space model), removal of the temporal delay in the joint velocity and position signals, and reduced mass of the delta robot arm linkages. Other identified constraints are currently being addressed in an exoskeleton system under development (see Discussion).

#### Force signal-processing

The noise (3 std.) in the force signal while the exoskeleton was turned on and holding a position was ±15 mN in x and y and ±0.2 mNm in yaw. Drift in the signal due to variable resistance in the slip-ring during yaw axis rotation was measured to be ±10 mN in x and y and ±0.1 mNm in yaw. Noise could not be reduced using a frequency filter because this would detriment the system bandwidth^23^. Therefore, a dead-band of ±25 mN in x and y and ±0.3 mNm in yaw was implemented on the force signal to ensure the noise and drift were not passed on to the admittance controller.

### Software programming

#### Exoskeleton

The servo-drivers were configured on initial set-up using motor parameter files provided by the supplier (Harmonic Drive LLC). Tuning of the current and velocity PID loops on the servo-drive and set up of control input and output functions (emulated quadrature encoder output, pulse width modulated (PWM) input, motor enable input) were performed within the software graphical user interface (GUI) provided by the supplier (HDM Software v71B24, Harmonic Drive LLC).

The force sensor calibration matrix, which converts raw analogue signals into 6DOF force values, was provided by the supplier (ATI Industrial Automation) and was implemented in the CompactRIO controller software (LabVIEW 2019, National Instruments). A force coordinate transform was used to convert the force sensor measurements from the force sensor coordinate frame to the mouse’s coordinate frame (**Supplementary Info. 6**). The CompactRIO controller executed all kinematics and admittance control calculations at 100 Hz cycle time. While all calculations were stored and executed on the CompactRIO controller, a GUI was available on a host PC that enable the experiment operator to adjust and monitor parameters during operation of the exoskeleton.

For complete software files for the CompactRio controller see the GitHub repository: https://github.com/bsbrl/exoskeleton

#### Imaging headstage

Data acquisition from the camera was configured using software GUI provided by the supplier (Vimba Viewer, Allied Vision). The main parameters were exposure time of 67 ms; gamma (contrast) of 2; Intensity of 50; Black level of 0; and pixel format of Mono12p. The gain was adjusted within the GUI for each mouse. The synchronization TTL signal generated by the CompactRIO controller was used to turn on and off the LED driver, which was used to synchronize the image series data with the exoskeleton data. Data were saved as an image series in .tiff format to a solid-state drive on the host PC.

#### Electrophysiology headstage

Digital data from the head stages and the synchronization signal generated by the CompactRIO controller were acquired on a USB interface board (RHD2000, INTAN) before being saved in binary format to a solid-state drive on the host PC. Data acquisition and pre-processing parameters were 20 or 30 kS/s (where S = samples) acquisition rate with 7.5 kHz antialiasing filter.

### Cranial Implants

The cranial implants for interfacing between the mouse and the robot were adapted from Ghanbari et al^8^. All cranial implant designs were 3D printed from photo curing polymer (Black v4 or Clear v1, FormLabs). The under-side of the cranial implants were designed to conform to the dorsal surface of the frontal bone plates, parietal bone plates, and occipital bone plate on an average mouse’s skull. Three 0-80 tapped holes were located on supporting structures extending outward from the implant and were used to chronically fix the cranial implant to the headpost. Two holes over the occipital plate were used to help secure the cranial implant to the skull using skull screws.

#### Imaging cranial implant

The cranial implant for imaging contained a 6 x 6 mm glass plug, adapted from (Hattori & Komiyama)^45^, that compressed a section of the dorsal cortex (-4 to +2 mm AP, -3 to +3 mm ML) into a single flat imaging plane. The glass plug was assembled from two pieces of glass. The top glass (Fused Quartz; MUHWA Scientific) was 0.5 mm thick and mated to reference faces on the cranial implant to define the orientation of the imaging plane. The bottom glass (Superfrost Plus Micro Slide; VWR) was 1 mm thick and was fixed to the underside of the top glass to define the depth of the imaging plane. The top and bottom glass pieces were glued together and to the cranial implant using Ultraviolet (UV)-curable optical glue (NOA81; Norland Product) and UV lamp (365nm, D11D; LIGHTFE). A plastic cap was attached to the headpost to protect the top glass from dust and debris.

#### Electrophysiology cranial implant

The cranial implant for electrophysiology experiments contained six probe entry ports that provided physical access for neural probes to brain (**Supplementary Info. 8**). The probe entry ports were each 1.4 mm diameter and were surrounded by a raft of support material that extended down 0.3 mm into the brain, through the craniotomy, to prevent dental cement from wicking up into the holes. The sides of the implant were raised by 1.2 mm to create a reservoir around the probe entry ports, which was filled with silicon elastomer (Kwik-Sil, World Precision Instruments) until just prior to experiments. A protective lid was secured over the silicon elastomer using miniature self-tapping screws (FF000CE094, JI Morris). The locations of the probe-entry ports were calculated to accommodate probe trajectories that intersected the primary motor and somatosensory cortices (0 mm AP; 1.5 mm ML; -2.37 mm DV), primary Barrel field (-1.8 mm AP; 3.5 mm ML; -3.5 mm DV), and primary visual cortex (-2.5 mm AP; 2.35 mm ML; -1 mm DV) in the left hemisphere, and the primary motor cortex (0.85 mm AP; -0.5 mm ML; -5 mm AV), anterior retrosplenial cortex (-1.05 mm AP; -0.5 mm ML; -2.7 mm DV), and posterior retrosplenial cortex (-3 mm BL; -0.75 mm ML; -1.5 mm DV) in the right hemisphere.

These probe trajectory calculations were performed using the kinematic description of the probe alignment mechanics^46^ and stereotaxic coordinates of the average mouse brain^47^. For ground connection, a skull screw (FF000CE094, JI Morris) with single strand 32 AWG wire soldered beneath the screw head was screwed to approximately 1mm depth into a pilot hole drilled through occipital bone plate. The solder on the skull screw was completely encased in dental cement once screwed in to avoid contact between exposed tissue and solder. The free end of the 32 AWG wire was then soldered to a gold-plated contact (36-122DKR-ND, Digikey) that was fixed to the titanium headpost.

### Surgery

All animal experiments were approved by the University of Minnesota Institutional Animal Care and Use Committee (IACUC). Mice were administered 2mg/kg of sustained-release buprenorphine (Buprenorphine SR-LAB; ZooPharm) and 2mg/kg of meloxicam for analgesia and inflammation prevention, respectively. Mice were anesthetized in an induction chamber containing 1–5% isoflurane in pure oxygen. The scalp was shaved and sterilized, followed by the application of sterile eye ointment (Puralube; Dechra Veterinary Products) to the eyes. Mice were then transferred and affixed to a standard rodent stereotax (Model 900LS; Kopf). The scalp above the dorsal cortex was excised using surgical scissors and the fascia was removed using a 0.5-mm micro curette (catalog no. 10080-05; Fine Science Tools). A stencil was aligned with bregma and lambda over the exposed skull and then guidelines were marked for the craniotomy and skull screws. Craniotomies and skull-screw pilot holes were performed manually using a high-speed dental drill. Once craniotomy skull flaps were removed, the exposed brain was immediately covered with a gauze pad soaked in sterile saline to keep the brain hydrated. The cranial implant was sterilized by soaking in 70% ethanol for 2min, followed by rinsing with sterile saline. The implant was gently placed on the skull and skull screws were inserted through the cranial implant into the two pilot holes in the occipital plate. The area of the skull surrounding the cranial implant was dried using cotton-tipped applicators, and then dental cement (Metabond, Parkell) was applied to adhere the implant to the skull. After the dental cement had fully cured, the headpost was attached to the cranial implant using three 0-80 screws. Exposed brain tissue was covered using silicon elastomer (Kwik-Sil, World Precision Instruments), and then a protective plastic lid was screwed in place (FF000CE094, JI Morris) over the silicon elastomer. Mice recovered on a heated recovery pad (catalog no. 72-0492; Harvard Apparatus) until fully ambulatory before returning to a clean home cage. Mice were subsequently allowed to recover for at least 14 days before commencement of training.

### Behavior

#### Oval track construction

The oval track arena contained an outer wall with two 27cm straight sections parallel to one another and connected by two semicircular sections 33 cm in diameter. The outer wall was 5 cm in height. The floor of the arena was removable and was fabricated from laser-cut acrylic sheet with a layer of 6 mm thick, textured neoprene rubber to provide grip for the mice. The exoskeleton allowed mice to walk along a trajectory that maintained a distance of 3 cm from the outer wall. Mice had control of their forwards direction through an admittance controller, with backwards direction disabled and the sideways and yaw directions controlled using a trajectory controller running in parallel to the admittance controller.

#### 8-maze construction

The 8-maze arena contained two circular goal arms with 36 cm diameter outer walls and 20 cm diameter inner walls, creating an 8 cm wide channel. The walls 3D printed from polylactic acid (PLA) and were covered with a layer of either white, black, or grey colored foam to provide visual contrast for the mice and were mounted to a baseplate that was laser-cut from acrylic sheet (**Supplementary Info. 7**). The floor of the 8 maze was covered with textured neoprene rubber. The two goal arms overlapped at the center of the 8-maze producing a zone where the mouse could decide on which goal arm to enter. This zone was named the turning zone and was triangle shaped with its narrowest point where the mouse entered and widest point where it connected to the beginnings of the two goal arms. A servo-controlled (SG90 9G, Miuzei) door at the end of the two goal arms blocked-off the route to the opposing goal arm, ensuring mice maintained a figure of 8 path through the maze. During initial behavioral training a second servo-controlled door (the training door) was located at the top of the turning zone to guide mice to enter the desired goal arm. The doors were actuated when an infrared break-beam (1528- 2526-ND, Digikey) located near the beginning of each goal arm was triggered by the mouse. When mice triggered a second infrared break-beam located at the end of each goal arm, sound (B07MPYWVGD, ARCELI via Amazon) and light (white LEDs) cues were activated that indicated to the mouse the correct goal arm to choose (**Supplementary Info. 7c**). When mice chose the wrong goal arm, a 3 second puff of air was released from ports in the goal arm walls via a solenoid valve (2W-025-08, Tailonz pneumatic) connected to a pressurized air line at 40 psi. When mice chose the correct goal arm, an 8 µL liquid reward was dispensed by gravity feed through a 1 mm inner diameter silicon tube via a solenoid valve (LHDA0533215H, The Lee Company). The silicon reward dispensing tube was mounted to a servo-actuator (SG90 9G, Miuzei) to allow height adjustment if necessary.

#### 8-maze implementation on the exoskeleton

When maneuvering the exoskeleton, mice started and finished the 8-maze on a disc treadmill located adjacent to the maze. The disc treadmill provided a point where the exoskeleton could be held stationary while neural recording probes were inserted and removed, or while imaging parameters were adjusted. Mice entered and exited the 8-maze along a trajectory that passed through the servo-controlled door at the end of the two goal arms, which was manually removed at these times (**Extended Data 5a-b**). Within the turning zone, mice had full control over their forwards/backwards, lateral, and yaw directions within position limits. Position limits at the global x and y boundaries of the turning zone were implemented in the exoskeleton controller by limiting the magnitude of the admittance controller output such that the resultant end position could not cross these boundaries (**Extended Data 5c**). A yaw axis limit was also implemented in the turning zone which scaled linearly from ±20 to ±45 degrees across the first 5 cm of the turning zone. Immediately prior to entering the turning zone, the mouse passed through a 1 cm wide by 3 cm long transition zone, within which the lateral and yaw velocities transitioned to 0 m/s and 0 rad/s, respectively, to ensure mice entered the turning zone moving in a purely forwards direction. In the goal arms, mice followed a circular trajectory that tracked along the center of the 8 cm wide channels. Here, mice had control of their forwards direction through an admittance controller, with backwards direction locked and the sideways and yaw directions controlled using a trajectory controller running in parallel to the admittance controller. The admittance controller output, which was a velocity vector for the mouse’s forwards direction, was always at a tangent to the circular trajectory, and so it acted to move the mouse off this trajectory. Therefore, a vector path correction calculation was implemented that terminated the admittance velocity vector at a point along the circular trajectory (**Extended Data 5d**). Trajectory control of the mouse’s lateral and yaw directions within the goal arms was calculated using vector notation (**Extended Data 5e**) which, in the case of the yaw axis, avoided the possibility of software errors from angle wrapping in trigonometric methods.

### Training

Training mice to perform the 8maze task on the exoskeleton involved five stages of training on different devices. Each device was designed to either acclimatize or teach mice one or more aspects of walking while head-fixed, controlling the exoskeleton, or performing alternating decisions within the 8maze.

#### Acclimatization

Mice were handled daily by the experimenter for 5 to 10 days before starting their training. Each time mice were introduced to a new training environment they were allowed to freely explore the environment for at least 20 minutes across 2 days to acclimatize them.

#### Head-fixed locomotion

The first stage of training involved head-fixation to a passive apparatus (without motors) which allowed mice to move through a physical environment under their own volition. Constraints on the motion of this apparatus were imposed that mimicked the exoskeleton implementation of the 8-maze.

#### Wheel training

The second stage of training was to teach mice to walk on a treadmill-style cylindrical treadmill while head-fixed. Mice were trained on this task daily until they walked or ran on the treadmill without struggling for 10 minutes, which took 10 days for all mice. The purpose of this training was to ensure mice would be comfortable when walking on the disc treadmill at the start and finish of the 8-maze. Videos were captured of this training to review performance and progress.

#### Exoskeleton turn training

The third stage of training was to teach mice to use the exoskeleton. To minimize stress, mice were initially held on the starting treadmill for 1 minute before entering the 8-maze and spent 5 minutes within the 8-maze. These times were increased up to 10 minutes each, as mice became acclimatized to the task. The training door was left in place for all training sessions, with the correct goal arm alternating between left and right each trial. Liquid reward and air puff punishment were not administered during training.

#### Freely behaving decision-making

The fourth stage of training was to introduce the decision element to the 8-maze and the associated reward and punishment for correct and incorrect decisions, respectively. Mice were trained on this task while freely behaving. For the first 5 sessions, the training door was left in place and an air puff was administered if mice went in the wrong direction around the 8-maze to reinforce the following of a figure of 8 path. If mice moved from one reward to the next without going in the wrong direction this was scored as a correct trial, whereas going in the wrong direction at least once between rewards was counted as an incorrect trial. Training sessions lasted either 15 minutes or until 35 correct trials were achieved. Once mice reached at least 80% proficiency, which took 2 to 5 days for all mice, they progressed to the decision-making task. For decision-making, the training-door was removed, and an air puff was administered if mice chose the wrong goal arm or if mice went in the wrong direction around the 8-maze.

#### Exoskeleton decision-making

The fifth and final stage of training was for mice to perform the decision-making task while maneuvering the exoskeleton. For each mouse, the training door was left in place until they were consistently licking from the reward port while maneuvering the exoskeleton and their turning proficiency was greater than 80%, which took 1 to 4 days for all mice. The training door was then left in place for the first 2 to 6 trials of each session to reinforce alternating-choice behavior (**Extended Data 7**).

### Controller optimization

#### Freely behaving velocity-acceleration profiles

A top-down view of mice (n = 3) exploring an open field arena was used to determine velocity and acceleration data for freely behaving mice^9^ (**Supplementary Video 1**). Markerless tracking^24^ was used to label points on the head, body, and tail-base. The mouse’s head yaw angle was calculated as the angle between a vector joining points on the head and the body center and a second vector joining the body center to the tail (**Extended Data 1a**). The mouse’s coordinate frame was defined with the origin at the body center and orientation aligned with the yaw angle. A coordinate transform from the global coordinate frame to the mouse coordinate frame was used to convert global velocity into the mouse’s velocity. Velocity and acceleration data were filtered (13-point median filter; 5-point mean filter) to reduce noise before calculating admittance profiles (**Extended Data 1)**.

#### Freely behaving pitch and snout height

A side-on view of mice (n = 4) locomoting along a straight section of the linear oval track was used to determine pitch and height data for freely behaving mice. Marker-less tracking^24^ was used to label points on the nose and ear, in addition to other body parts (**Extended Data 2**). Mouse velocity was calculated from the average velocity of the nose and ear points. Pitch angle was calculated as the angle between a vector joining the nose and ear points and the global horizontal axis.

#### X admittance tuning

The oval track arena was implemented on the exoskeleton using the same combined admittance-trajectory control approach as the 8-maze arms (**Extended Data 5d-e**). The range of mass and damping values to test were estimated from preliminary force data collected from stationary, head-fixed mice and from velocity and acceleration data collected from markerless tracking of freely behaving mice. The 14 combinations of mass and damping values were tested in 4 mice across 20 sessions, with 1 to 2 combinations tested each session (mouse 1 = 2 sessions, 2 combinations; mouse 2 = 3 sessions, 3 combinations; mouse 3 = 9 sessions, 11 combinations; mouse 4 = 6 sessions, 12 combinations) (**Extended Data 2**).

#### Y and Yaw tuning

The range of mass and damping values to test were estimated from preliminary force data collected from stationary, head-fixed mice and from velocity and acceleration data collected from markerless tracking of freely behaving mice. The 13 different mass and damping combinations in y (7) and yaw (6) were tested in 2 mice across 15 sessions, with 1 to 3 combinations each session (mouse 1 = 15 sessions, 13 combinations; mouse 2 = 14 sessions, 10 combinations) (**Extended Data 2**).

#### Gait analysis

A side-on view of mice (n = 3) locomoting along a straight section of the linear oval track was used for gait analysis of freely behaving mice and mice maneuvering the exoskeleton. Marker- less tracking^24^ was used to label points on the nose, left ear, left forepaw, and left hind paw, in addition to other body parts. Mouse velocity was calculated from the average velocity of the nose and ear points. The start and end of individual steps were extracted from the data by identifying times where paws were stationary (velocity of 0 cm/s). Jitter artefacts in the labelling were removed by filtering data (median filter, 5 points) and using exclusion criteria (step size < 1cm or > 10 cm, step duration < 0.05 s).

### Data Recording

#### Exoskeleton

Up to 31 variables were saved from the CompactRIO controller during each experiment with temporal resolution of 10 ms (100 Hz cycle time). These variables and their units were the actual controller cycle time (ms); joint positions (radians); operational-space positions (m, rad); forces generated in the mouse’s frame (N, Nm); target velocity output from admittance controller in the mouse’s frame (m, rad); duty cycle of PWM velocity command sent to the servo-drivers (%); and admittance controller mass (kg, kg.m^2^) and damping values (Ns/m, Nm.s/rad). In 8maze behavioral experiments, the digital signals controlling sound cues, air puff, reward, and data synchronization were also saved.

#### Electrophysiology

One week prior to experiments, mice were anaesthetized using isoflurane and the silicon elastomer was replaced with a silicone gel (Dowsil 3-4680, Dow Chemical Company) that was compliant enough for neural probes to travel through. A protective cap that contained probe entry ports was also fixed to the cranial implant. On the day of recording, neural probes were aligned to a dummy cranial implant so that only the depth axis was needed during probe insertion on the mouse subject. The probes were dyed with a 0.1% weight/volume solution of red fluorescent dye (CellTraker^TM^ CM-Dil, ThermoFisher Scientific) in ethanol by passing the probe through a 2 to 5 μL drop of solution at the end of a pipette. In the event an entry port was for two recording sessions, a green fluorescent dye was used (CellTracker^TM^ Green CMFDA, ThermoFisher Scientific) in the second session. At least 2 hours prior to recording, mice were anaesthetized so that a ground wire could be soldered to the ground contact on the headpost. The ground wire was a single core 32 AWG wire that extended 60 mm off the back of the headpost. The linear axis of the probe alignment mechanics was used to insert the probes into the brain at a rate of 5 to 50 μm/s using the software provided by the manufacturer (Pathway^TM^, New Scale Technology). Probe insertion took on average 7 ± 2 minutes. At the end of recording, probes were removed at 250 μm/s. The probes were then immediately rinsed with a stream of de-ionized water, then allowed to soak in tergazyme for a minimum of 2 hours, and finally soaked in de-ionized water for a minimum of 1 hour.

#### Imaging

Immediately prior to recording, mice were head-fixed to the exoskeleton while free to run on a disk treadmill. If necessary, the imaging glass was cleaned of dust and debris using cotton tipped swabs soaked in 70% ethyl alcohol. The gain was manually adjusted while viewing a live feed of the cellular fluorescence in the camera GUI (Vimba Viewer, Allied Vision). Imaging data were acquired while mice were learning to use the exoskeleton (n = 4 mice; 2 to 9 sessions per mouse). Light sources in the 8-maze were turned off to reduce noise.

### Data analyses

#### Exoskeleton

Mouse velocity and acceleration when maneuvering the exoskeleton were calculated by differentiating exoskeleton position with respect to time and then filtering to reduce noise (13- point median filter; 5-point mean filter). Force and position data were unfiltered. Admittance planes of the controller, for visualization in figures, were manually fit to velocity (v) – acceleration (a) – force (F) data by adjusting mass (m) and damping (c) values within the equation F = ma + cv.

#### Imaging

Image registration and detection of neurons was performed in the open-source software suite2p using the default settings for 1-photon imaging and cell diameter of 5 pixels^29^. Fluorescence traces from each cell (Fcell) and neuropil (Fneu) were band pass filtered (4^th^ order Butterworth, 0.05 to 5 Hz passband) and then used to calculate the change in fluorescence (ΔF/F) using the equation ΔF/F = (Fcell – 0.7*Fneu)/mean(Fcell – 0.7*Fneu). Cells were removed if they did not meet several inclusion criteria on the ΔF/F signal (3*standard deviation < maximum; 1*standard deviation < 50%; maximum between 20% and 500%) and cell the morphology (number of pixels in cell soma between 20 and 400 pixels; soma aspect ratio between 0.7 and 1.4). Brain atlas boundaries were scaled up from the source resolution (10 um) to the imaging resolution (3.45 um) and then the two were manually aligned using a transform (translation and rotation). Before linear regression, mouse location data from the exoskeleton (100 S/s) was resampled at the imaging sample rate (15 S/s).

The kernel (K; location bin x cell) of Beta weights was generated from the inferred spike rates (Y; time x cell) and a binary predictor matrix (X; time x position bin) of the mouses location in the 8-maze by rearranging the equation Y = X*K to solve for K^38^. Linear regression using the Moore-Penrose pseudo-inverse was determined to be suitable for this because the mouse could only occupy 1 location bin at a time, resulting in the predictor matrix having full rank. The predictor matrix was generated by binning the turning zone and arms into approximately 2 cm length bins and then assigning the column of the predictor matrix a value of 1 when the mouse was present in the corresponding location bin. Inferred spike rates were output directly from suite2p. The inferred spike rates when the mouse was stationary was zeroed before linear regression to avoid fitting to quiescent states, and kernel events with less than 5 instances were zeroed to avoid over-fitting to sparse events. Each cell’s Beta weights in the resultant kernel can be interpreted as the average response of that cell to the location bins in the 8-maze. To compare the response across cells, each column (cell) of the Kernel was normalized and then the columns were arranged in order of the location onset of their maximum. Note that for visualization the kernel is transposed in **Fig. 4** and **Extended Data 8.**

#### Electrophysiology

The mean of all the voltages across channels that were visually determined to be within the brain (based on low noise amplitude) were subtracted from all channels to reduce the overall noise. The noise-reduced data were then processed using Kilosort spike sorting software (Kilosort 2.0^30^) and manually curated using the ‘phy’ GUI^31^. During post-processing, neural data was bandpass filtered (4^th^ order Butterworth, 4 to 250 Hz passband) before calculating mean waveforms, spike amplitudes, and noise amplitudes. To calculate spike rates, the number of spikes were summed within a 50 ms length window, run across the neural data in 20ms increments, which acted to resample the data at 50 S/s. Spike rates were filtered by convolving a Guassian pulse with 260 ms width and 20 ms standard deviation, and then z-scored. Before linear regression, mouse position data from the exoskeleton (100 S/s) was resampled at the sample rate of the z-scored spike rate data (50 S/s).

The kernel (K; location bin x cell) of Beta weights was generated from Guassian smoothed z- scored spike rates (Y; time x cell) and a binary predictor matrix (X; time x event) by solving the equation Y = X*K using reduced rank regression^48^. Reduced Rank Regression was used here because multiple events could coincide with one another within the navigational decision-making task. Spatial location events in the predictor matrix were generated by binning the turning zone and arms into approximately 1 cm length bins and then assigning the column of the predictor matrix a value of 1 when the mouse was present in the corresponding location bin. In addition to location, diagonalized binary events were defined for auditory cues and reward dispensing using the method described in Steinmetz et al^2^. Lastly, several continuous variables were z-scored and included in the predictor matrix (velocity, acceleration and force in x, y, and yaw). Kernel spatial location events with less than 5 instances were zeroed to avoid over-fitting. Each cell’s Beta weights across the spatial location events in the resultant kernel can be interpreted as the average response of that cell to the location bins in the 8-maze. To compare the response across cells, each column (cell) of the Kernel was normalized and then the columns were arranged in order of the location onset of their maximum. Note that for visualization the kernel is transposed in **Fig. 5 and Extended Data 9**.

### Statistical Analyses

#### Admittance tuning

Velocity, acceleration, and force data are continuous variables. Therefore, to ensure each sample was an independent observation, we only included the peaks in our analysis. Box and whisker plots in **Fig. 2f** and **Extended Data 3e** show the distributions of all positive peaks (n values included in **Extended Data 3e**). To compare the velocity and acceleration peak distributions across different admittance controller settings, peaks were resampled to n = 100 peaks to avoid exaggerating differences in the data from oversampling. Statistical parameters for **Fig. 2f** are as follows: Velocity: 1-way ANOVA, F range = 0.3 to 0.5, n = 100 samples per group, within-groups df = 1. Acceleration: 1-way ANOVA, F range = 0.07 to 0.4, n = 100 samples per group, within-groups df = 1. Force: 1-way ANOVA, F = 0.12, n = 100 samples per group, within-groups df = 1).

#### Gait

Steps were only included in analysis if the entire mouse was visible in frame. Statistical parameters for **Fig. 2h** are as follows: Forepaw step length: 1-way ANOVA, F = 1.2, n = 121 steps freely behaving, n = 268 steps maneuvering the exoskeleton, within-groups df = 1. Hind paw step length: 1-way ANOVA, F = 0.23, n = 136 steps freely behaving, n = 235 steps maneuvering the exoskeleton, within-groups df = 1. Forepaw cadence: 1-way ANOVA, F = 27, n = 67 steps freely behaving, n = 198 maneuvering the exoskeleton, within groups df = 1. Hind paw cadence: 1-way ANOVA, F = 52, n = 82 steps freely behaving, n = 165 steps maneuvering the exoskeleton within groups df = 1.

#### 8-maze performance

Success rate used to evaluate turning proficiency (**Fig. 3d**) and decision making (**Fig. 3h**) are binary metrics (1 = correct trial; 0 = incorrect trial). Therefore, mean and 95% confidence intervals were calculated using the Clopper-Pearson binomial (MATLAB function *binofit*). Linear trends in data were determined using the least-squares approach. Comparison in decision making between freely behaving and exoskeleton maneuvering groups were calculated using Kruskal-Wallis test because it does not assume a normal distribution in the data. For this test, the 1^st^ session was excluded from both groups because mouse performance was consistently lower in both freely behaving mice and mice maneuvering the exoskeleton, which was thought to be caused by the change in task environment from freely behaving training (training door in place) to freely behaving decision making, or from exoskeleton training (training door in place) to exoskeleton decision-making. Statistical parameters for **Fig. 3h** are as follows: Kruskal-Wallis test, n = 1609 trials freely behaving, n = 361 trials maneuvering the exoskeleton, Chi-squared = 2.0, within groups df = 1)

## ACKNOWLEDGEMENTS

We acknowledge the staff at Research Animal Resources, University of Minnesota, for animal care and housing; Jacob O’Brien for his initial exploration of the exoskeleton concept; Daniel Surinach for assistance training models in DeepLabCut; Jia Hu for assistance using Suite2p and processing ΔF/F traces; Emily Troester for assistance with code for video acquisition; Arun Cherkkil for consultation on mesoscale imaging headstage components. Funding from the UMN Mechanical Engineering Department, MnDRIVE RSAM, The McKnight Foundation, the Minnesota Robotics Institute (MnRI), National Institutes of Health (NIH) grant 1R01NS11128, and BRAIN Initiative grants RF1NS113287 and RF1NS126044 are gratefully acknowledged.

## AUTHOR CONTRIBUTIONS

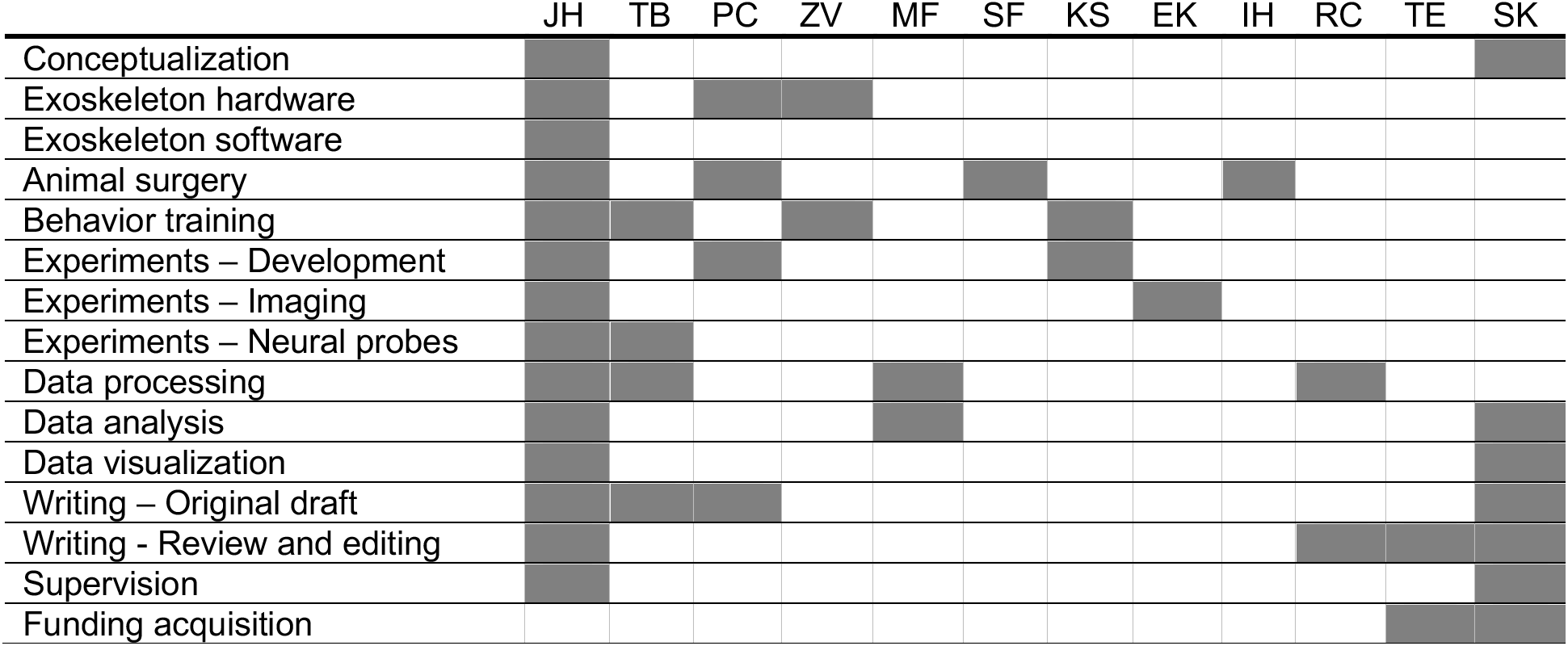

## DATA AVAILABILITY

The data sets reported here are openly available in the GitHub repository: https://github.com/bsbrl/exoskeleton

## CODE AVAILABILITY

All code for controlling the robot and for data analysis is available in the GitHub repository: https://github.com/bsbrl/exoskeleton

## LIST OF SUPPLEMENTARY MATERIALS

**1) Extended data figures:** Document consisting of nine captioned figures containing data and schematics that support the results and methods presented in the main text.

**2) Supplementary Info:** Document consisting of eight captioned figures containing data and schematics that support the results and methods presented in the main text.

**3) Supplementary software and Supplementary CAD files:** Can be found at GitHub Repository: https://github.com/bsbrl/exoskeleton

**4) Supplementary Videos:** Five captioned videos containing video data and schematics that support the results and methods presented in the main text.

Supplementary Video 1: Velocity-acceleration profiles of freely behaving mice and mice maneuvering the exoskeleton during tuning of the x axis.

Video (8x speed) showing top-down view of a mouse in open field arena with several body points labelled using labelled using DeepLabCut markerless tracking software (left window), and the corresponding velocity-acceleration profile for forwards/backwards motion in the mouse’s frame of reference (grey; right window). Transition at half-way to a mouse maneuvering the exoskeleton around the linear oval track with tuned mass and damping values and the corresponding velocity-acceleration profile (blue).

Supplementary Video 2: Gait dynamics of freely behaving mice and mice maneuvering the exoskeleton.

Video (1x speed) showing a side-on-vide of a freely behaving mouse locomoting along a 27 cm long straight section of the linear oval track (left window), with left paws and other body points labelled using DeepLabCut marker-less tracking software, and the corresponding gait metrics for steps in the video (large grey dots) overlaid on top of the gait metrics for all mice (right window). Transition at half-way to the same mouse maneuvering the exoskeleton with metrics for steps in the video (large blue dots) overlaid on top of the gait metrics for all mice.

Supplementary Video 3: Mouse performing the navigational decision-making task while maneuvering the exoskeleton.

Video (2x speed) showing a trained mouse performing the navigational decision-making task (sensory-cue guided alternating choice) while maneuvering the exoskeleton. Top half of the video shows (left) view from behavioral camera mounted to the headstage and (top-right and bottom-right) views from 2 cameras placed around the behavioral arena. The bottom half of the video shows (top panels) the force time series in x, y and yaw, and (bottom panels) the velocity and acceleration profiles in x, y and yaw, where thick lines indicate the mouse is in the turning zone.

Supplementary Video 4: Mescoscale cellular resolution imaging of the dorsal cortex of a mouse navigating the 8-maze arena

Video (8x speed) showing a mouse completing 16 turns in the 8-maze arena while maneuvering the imaging headstage on the exoskeleton (top-left window), with corresponding video of fluorescence in the mouse’s cortex captured using the imaging headstage (bottom-left window).

Single cell activity is visible in the four, 1x1 mm, regions of interest (ROIs) (right windows). Transition half-way to a maximum intensity image of the brain (bottom-left window), and to plot of the ΔF/F traces from all cells in each ROI (right windows).

Supplementary Video 5: Multi-site electrophysiology recordings in a mouse performing the navigational decision-making task in the 8-maze arena

Video (10x speed) showing a mouse on the disk treadmill (top-left window) as probes are inserted through the cranial implant (bottom-left window) into the two anterior probe insertion sites (top-right window). Transition at half-way to video (2x speed) of the mouse completing a correct right turn decision (top-left window) with a corresponding plot of voltage signal traces from 23 channels across the 2 neural probes showing spiking signals, where colored bar indicates the probe (bottom left window; dark blue, M1; light blue, M1/SS). In the right window, the mean waveforms of 24 cells identified on the 2 probes are shown (thick colored lines; dark blue, M1; light blue, M1/SS) overlaid with the spiking activity of these cells (thin black line) as the spikes occur.

## Extended Data

**Extended Data Fig. 1:**
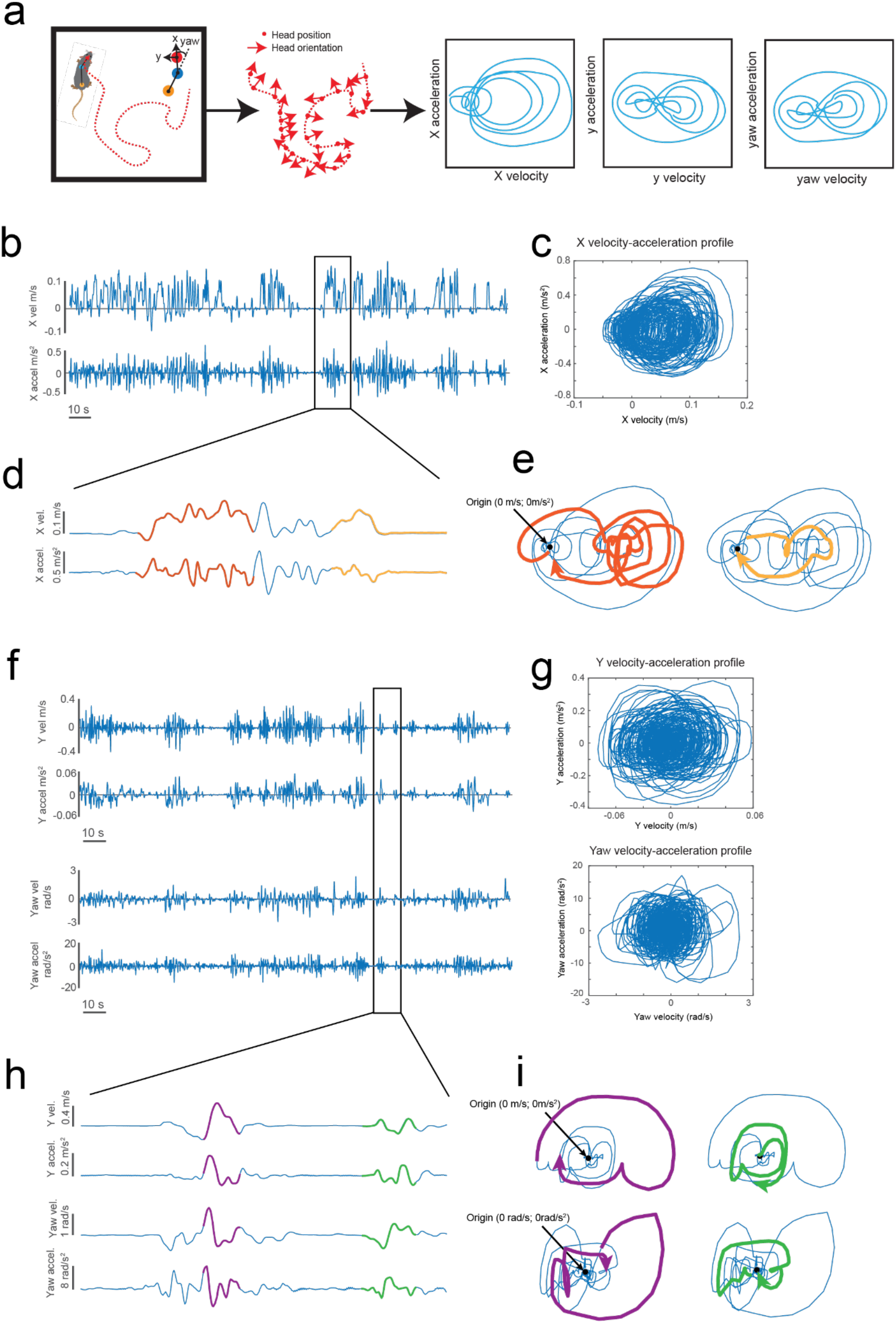
Velocity – acceleration profiles of freely behaving mice (a) Schematic of the method used to calculate velocities and accelerations in the mouse’s coordinate frame from marker-less tracking data. (b) Example data of velocity and acceleration time series in the mouse’s x axis (forwards/backwards). (c) Velocity-acceleration profile of the data in (b). (d) Close-up view of a 20 s clip (blue trace) of the data in (b), with two periods of motion highlighted (orange and yellow traces). (e) Velocity-acceleration profiles of the data in (d). (f) Example data of velocity and acceleration time series in the mouse’s y (lateral) and yaw (rotation left and right) axes (top, y; bottom, yaw). (g) Velocity-acceleration profiles of the data in (f), (top, y; bottom, yaw). (h) Close-up view of a 15 s clip of the data in (f), with two periods of motion highlighted (purple and green traces). (i) Velocity-acceleration profile of the data in (h).

**Extended Data Fig. 2:**
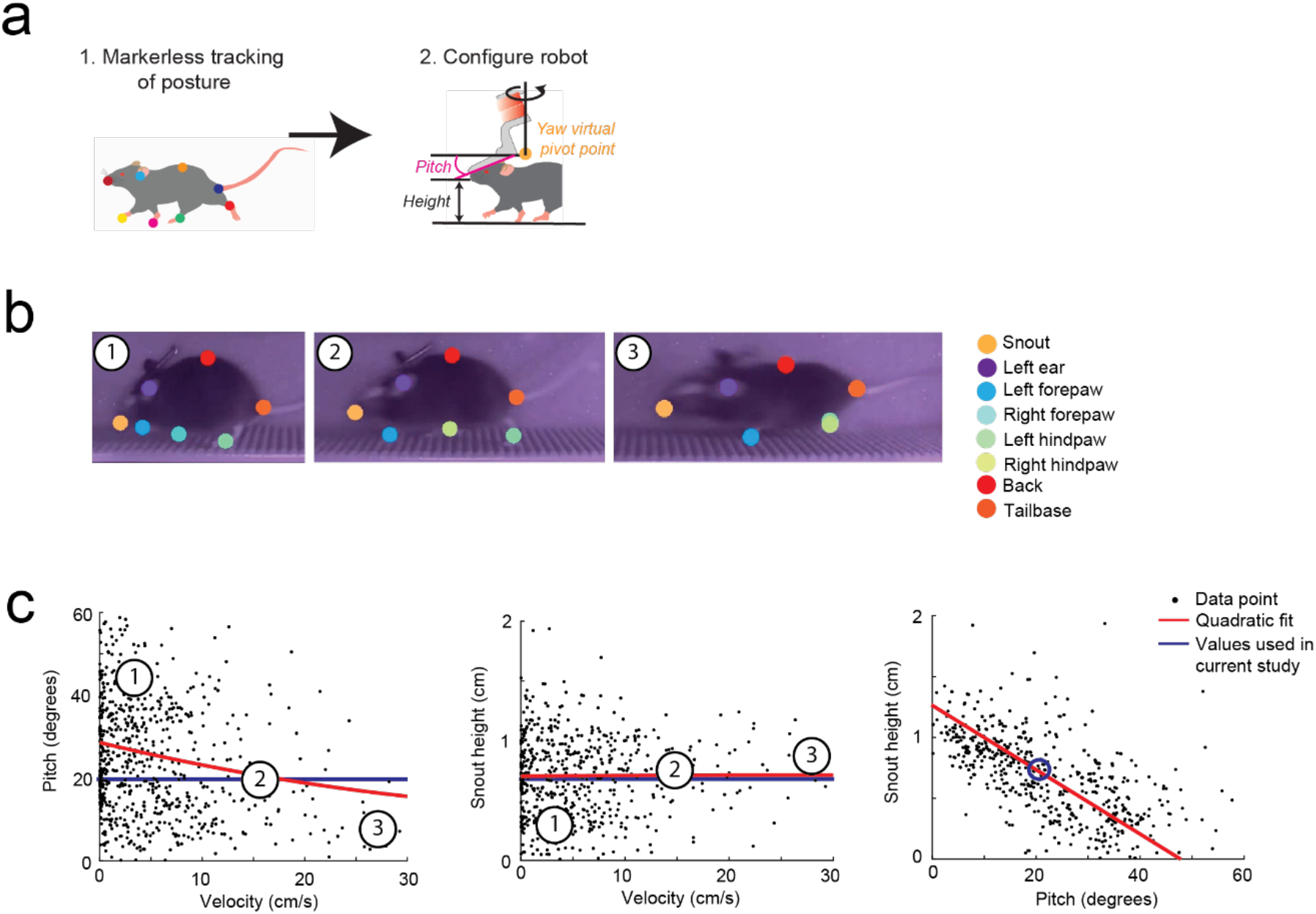
Pitch and height configuration for the exoskeleton (a) Schematic of the methodology used to configure the pitch and height settings of the exoskeleton. (b) Images of a freely behaving mouse overlaid with digitally labelled points on the left paws and other body parts, moving at 3 different forwards velocities (1., low velocity; 2., medium velocity; 3., high velocity). (c) Plot of head pitch angles and velocity for all mice (n = 4), with a quadratic fit to the data (red) and the pitch value implemented on the exoskeleton (blue). Numbers correspond to images in (b). (d) Plot of snout height and velocity for all mice (n = 4), with a quadratic fit to the data (red) and the height value implemented on the exoskeleton (blue). Numbers correspond to images in (b). (e) Plot of the pitch angle and snout height for all mice (n = 4), with a quadratic fit to the data (red) and the height value implemented on the exoskeleton (blue).

**Extended Data Fig. 3:**
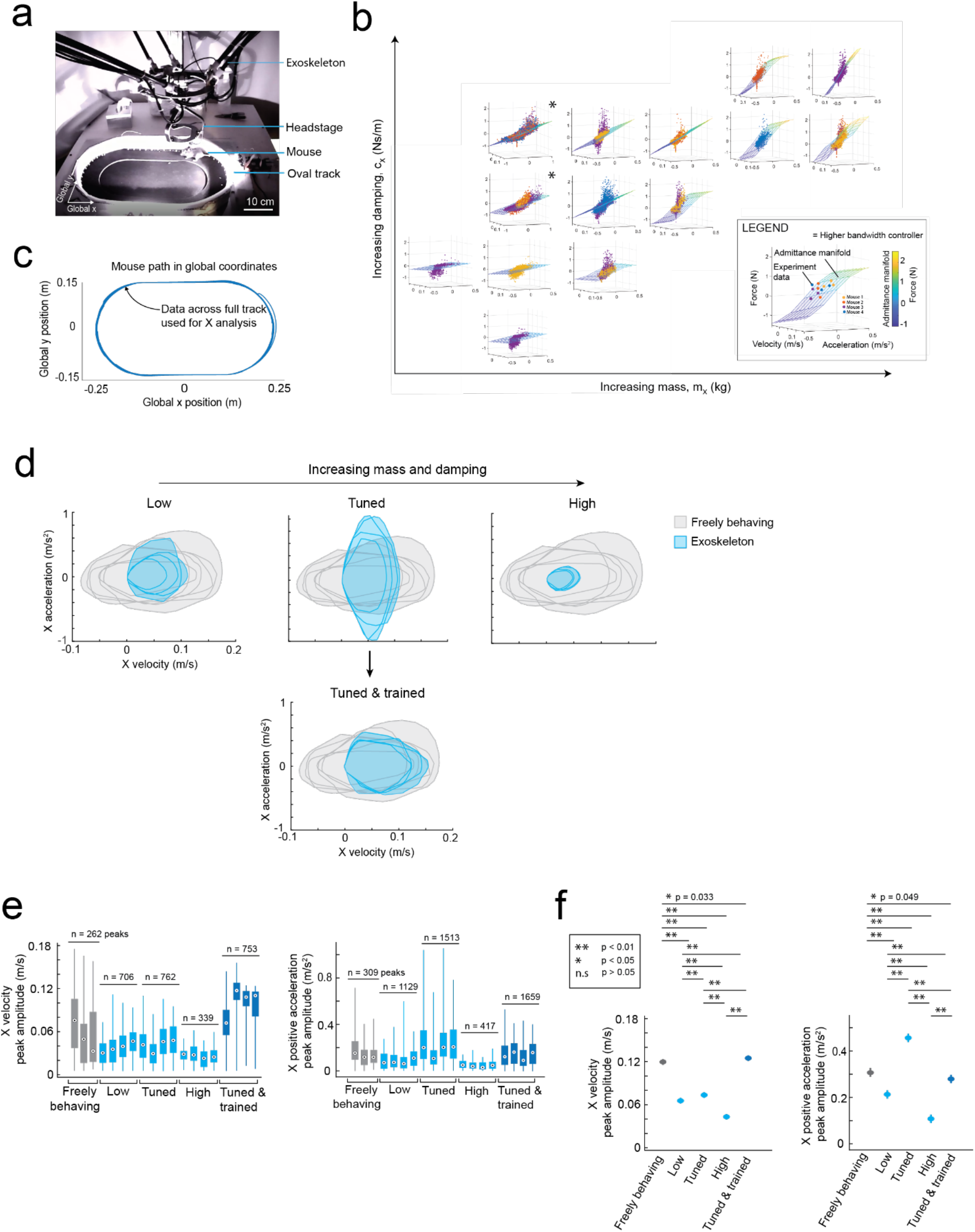
Tuning the admittance controller in the mouse’s x axis (a) Image of a mouse maneuvering the exoskeleton around the linear oval track used in experiments for tuning the admittance controller in the mouse’s x axis. (b) Plots of raw velocity-acceleration-force data from all mice (n = 4) and the associated admittance plane of the controller, arranged in order of increasing mass (left to right) and damping (bottom to top) for the 14 combinations of values evaluated. (Inset) Legend showing example plot with labelled axes and color-scale of the admittance plane (or manifold). (c) Example data of the mouse’s path around the oval track, where all data within the oval track was used to evaluate admittance tuning. (d) Data bounds of the velocity-acceleration profiles of freely behaving mice (grey; 3 mice; 6 sessions) and of mice maneuvering the exoskeleton with low (top-left), tuned (top-center), and high (top-right) mass and damping values (blue; 4 mice; 4 sessions), and with tuned mass and damping values after training (bottom-center; blue; 1 mouse; 4 sessions). (e) Distribution of the velocity peak amplitudes (left) and of acceleration peak amplitudes (right) of individual mice, when freely behaving, and when maneuvering the exoskeleton with low, tuned, and high mass and damping values, and tuned mass and damping values after training. (f) Mean and 95% confidence intervals of all velocity (left) and acceleration (right) peaks, with p-values calculated using 1-way ANOVA with n = 100 representative sample peaks per group.

**Extended Data Fig. 4:**
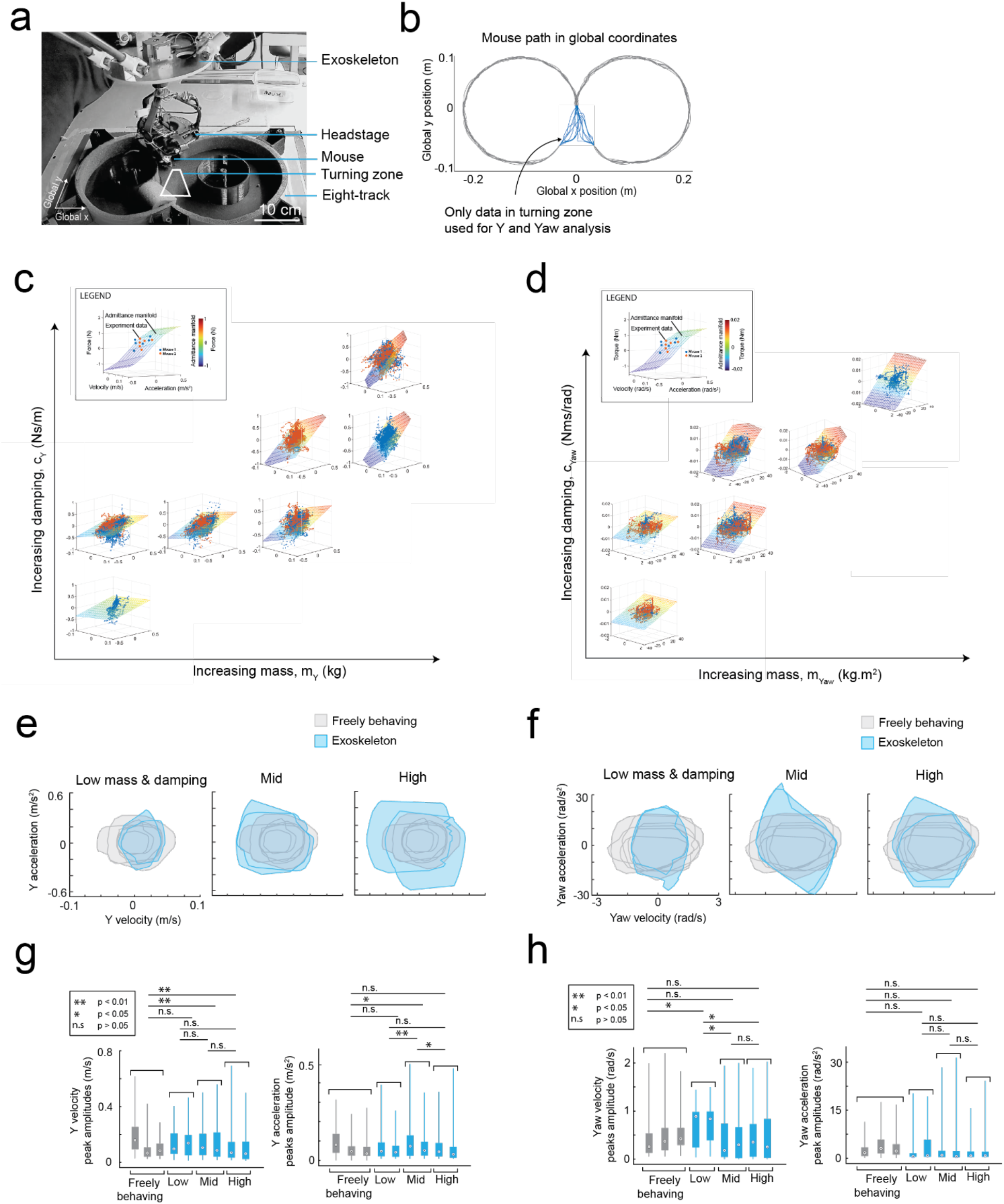
Tuning the admittance controller in the mouse’s y and yaw axes (a) Image of a mouse maneuvering the exoskeleton locomoting around the 8-maze arena used in experiments for tuning the admittance controller in the mouse’s y and yaw axes. (b) Example data of the mouse’s path around the 8-maze arena, showing the turning zone in the center of the arena which was used to evaluate admittance tuning. (c) Plots of raw data velocity-acceleration-force data in the mouse’s y axis for all mice (n = 2) and the associated admittance plane of the controller, arranged in order of increasing mass (left to right) and damping (bottom to top) for the 7 combinations of values evaluated. Legend (inset) showing example plot with labelled axes and color-scale of the admittance plane (or manifold). (d) As in (c) but for the 6 combinations of mass and damping evaluated in the mouse’s yaw axis. (e) Data bounds of velocity-acceleration profiles in the mouse’s y-axis for freely behaving mice (grey; 3 mice; 6 sessions) and for mice maneuvering the exoskeleton with low (left), medium (center), and high (right) mass and damping values (blue; 2 mice; 2 sessions). (f) As in (e) but for the mouse’s yaw axis. (g) Distribution of the velocity peak amplitudes (left) and of acceleration peak amplitudes (right) of individual mice, when freely behaving, and when maneuvering the exoskeleton with low, tuned, and high mass and damping values. Significance (p-values) calculated using 1-way ANOVA between groups. (h) As in (g) but for the mouse’s yaw axis.

**Extended Data Fig. 5:**
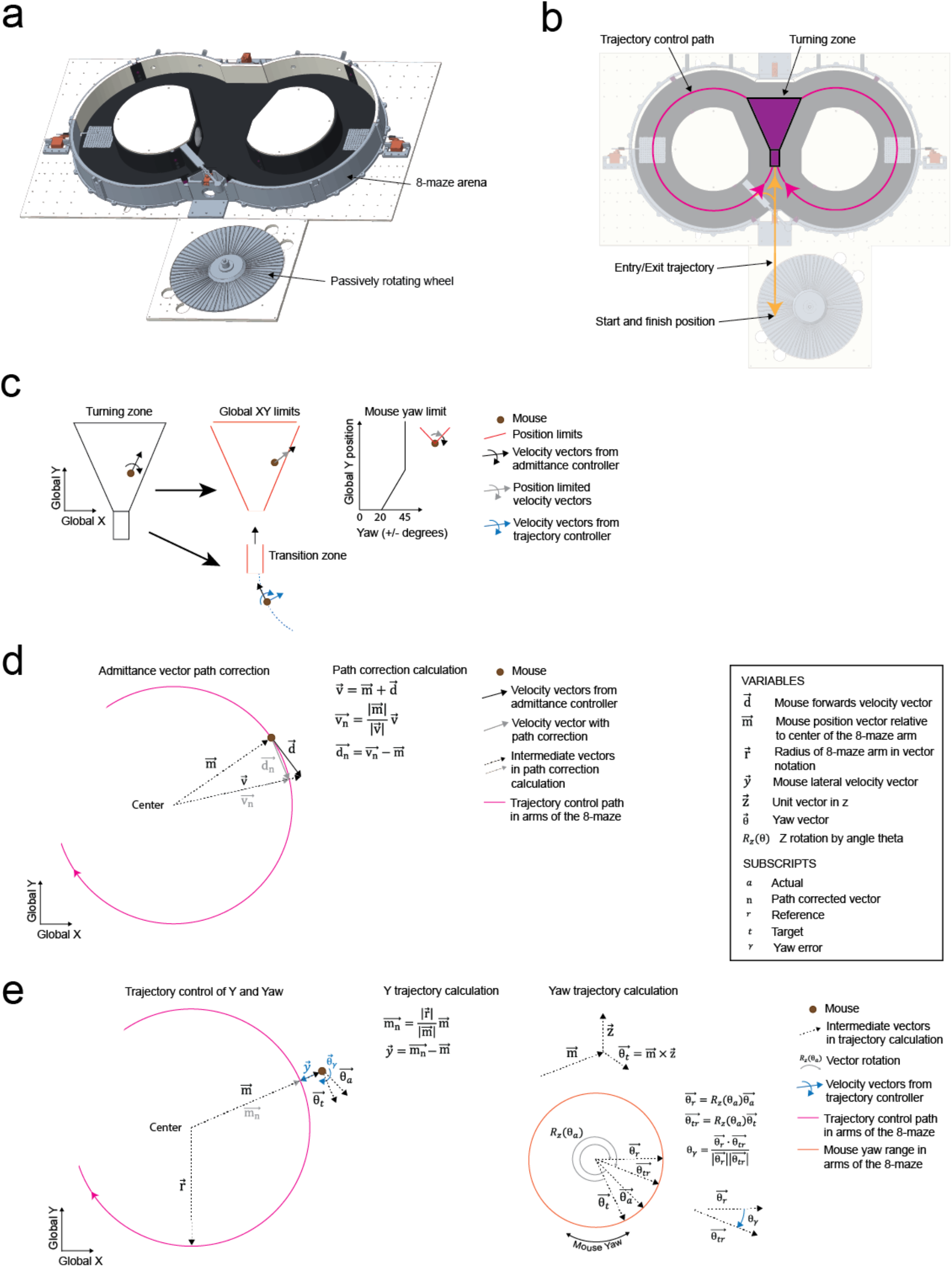
Implementation of the 8-maze arena on the exoskeleton controller (a) CAD rendering of the 8-maze arena and the adjacent passively rotating wheel. (b) Top-down view of the 8-maze arena and the adjacent wheel, overlaid with arrows indicating the trajectory control path through the left and right arms of the maze (magenta), trajectory control path for entry and exit into the maze (green), and the turning zone in the center of the maze (purple box with black outline). (c) Implementation of the turning zone, where global position limits (red) prevented the mouse from entering or exiting the zone except at defined points to avoid collisions of the exoskeleton with the arena walls. The mouse’s yaw axis was also limited to ± 45 degrees to prevent the mouse from attempting to turn around. Within these limits, the mouse had full control of its x, y, and yaw axes. (d) Vector path correction used in the arms of the 8-maze to ensure the mouse’s forwards velocity vector (d) was constrained to trajectory control path. (e) Implementation of trajectory control in the mouse’s y (y) and yaw (θ) axes within the arms of the 8-maze. To avoid angle wrapping, the mouse’s yaw angle was coordinate transformed (Rz) by the global yaw angle (8a) before calculating the yaw angle error (8ψ).

**Extended Data Fig. 6:**
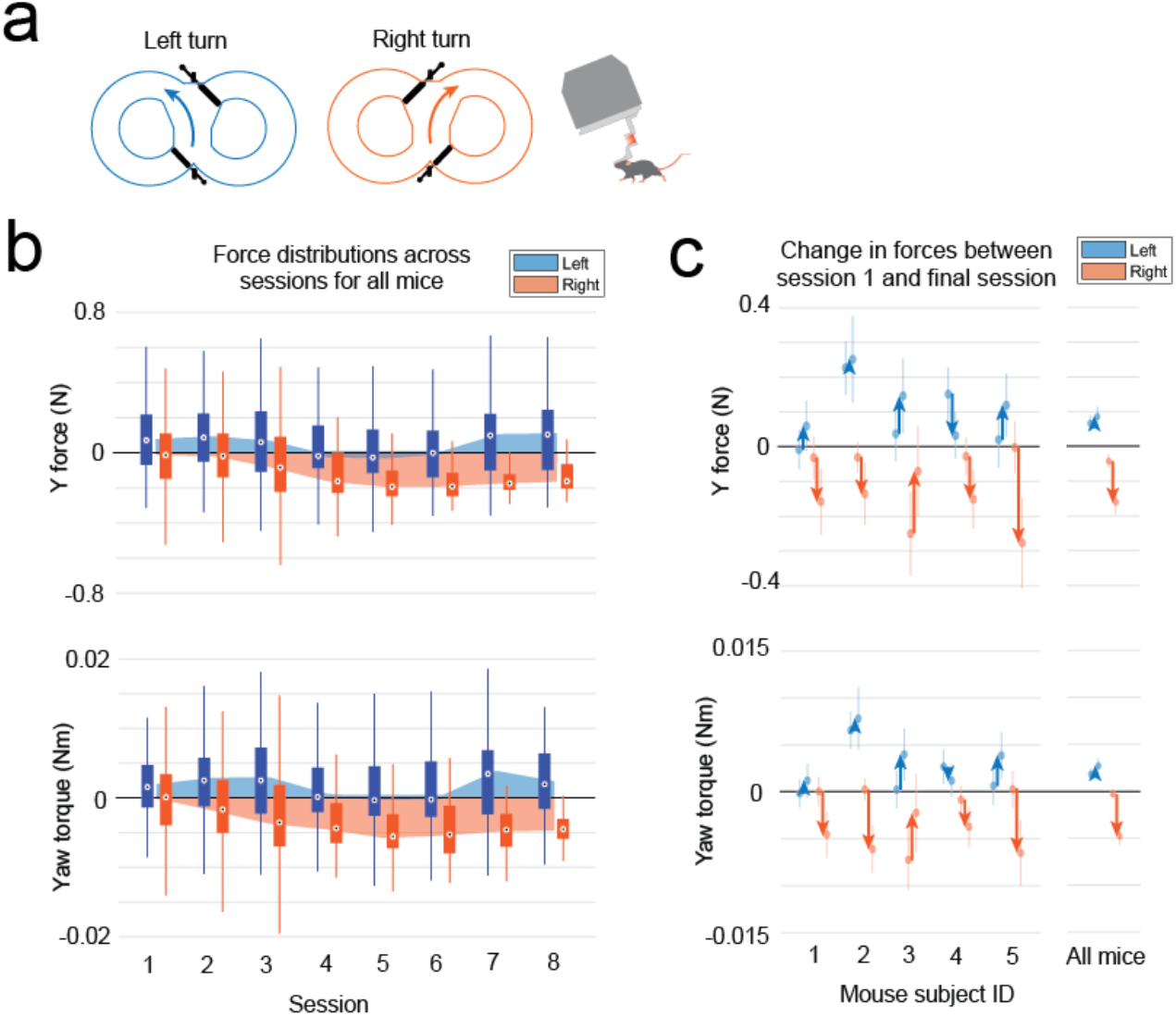
Forces during exoskeleton turn-training in the 8-maze arena (a) Color coded symbols of the 8-maze arena with both doors in place (turn-training) and mouse maneuvering the exoskeleton indicating the data in the figure. (b) Distribution of force peaks during left (blue) and right (orange) turns through the turning zone of the 8maze for all mice (n = 5) across 8 sessions (top, y forces; bottom, yaw torques). (c) Change (arrows) in force peaks for each mouse (left) and for all mice (right) between their first and final sessions for left (blue) and right (orange) turns (top, y forces; bottom, yaw torques).

**Extended Data Fig. 7:**
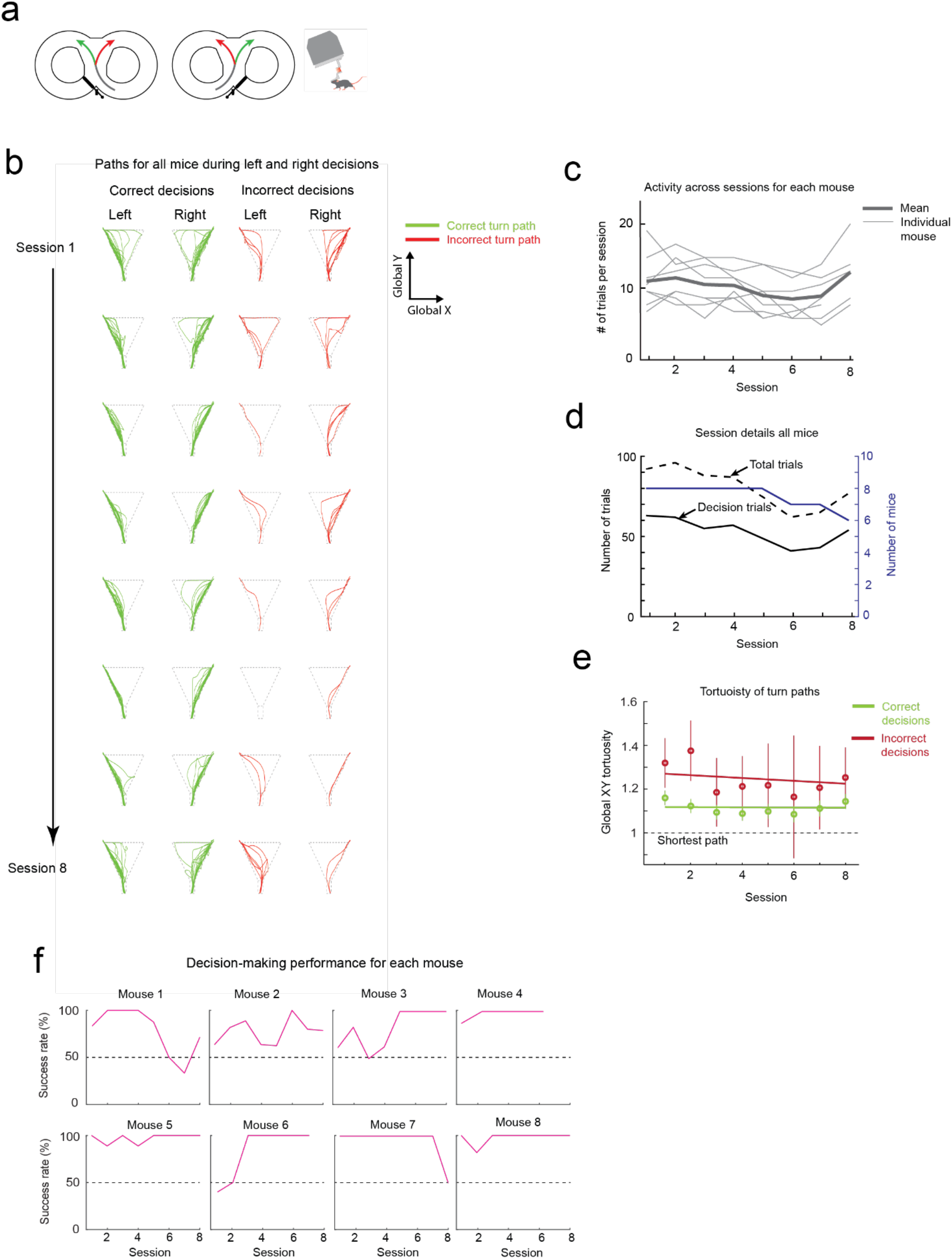
Performance during the decision-making task on the exoskeleton (a) Color coded symbols of the 8-maze arena with one door (decision-making) in place and mouse maneuvering the exoskeleton indicating the data in the figure. (b) Paths through the turning zone for correct (green) and incorrect (red) trials for all mice (n = 8) across 8 sessions. (c) The total number of trials per session for individual mice and the mean across all mice (n = 8). (d) Plot of the total number of trials in each session (left-axis; dashed black line) and the number of decision trials in each session (left-axis; solid black line) where decision trials are less than total trials because mice had 2 or more trials at the start of each session with the training door in place in the 8-maze arena. The number of mice in each session (right-axis; blue line). (e) Mean and 95% confidence intervals of the path tortuosity during decision trials for all mice (n= 8; green, correct decisions; red, incorrect decisions). (f) Decision-making performance for each mouse while maneuvering the exoskeleton.

**Extended Data Fig. 8:**
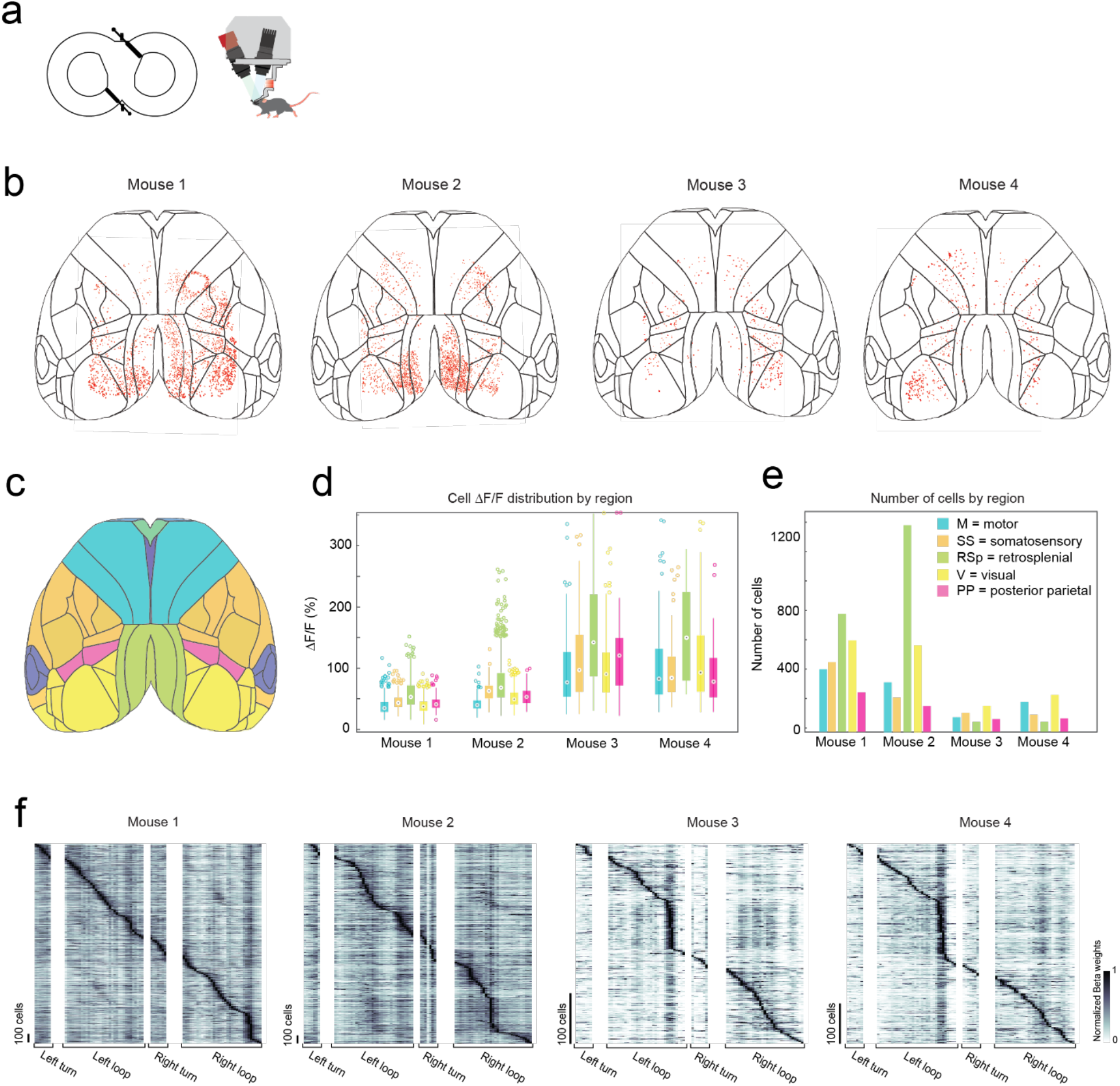
Summary of mesoscale imaging results across all mice (a) Symbols of the 8-maze arena with both doors in place (turn-training) and mouse maneuvering the mesoscale imaging headstage indicating the data in the figure. (b) Anatomical locations of all cells imaged in each mouse during an example session in the 8- maze arena. (c) Atlas of the mouse cortex with color coded functional regions. (d) Fluorescence (ΔF/F) distribution for each mouse broken down by brain region. (e) Number of cells imaged in each mouse broken down by region. (f) Kernel matrix of normalized Beta weights for each mouse, obtained using linear regression on the inferred spike rate of each cell and the location of the mouse in the 8-maze arena (82 location bins), with rows in the kernel sorted by onset of the maximum beta weight.

**Extended Data Fig. 9:**
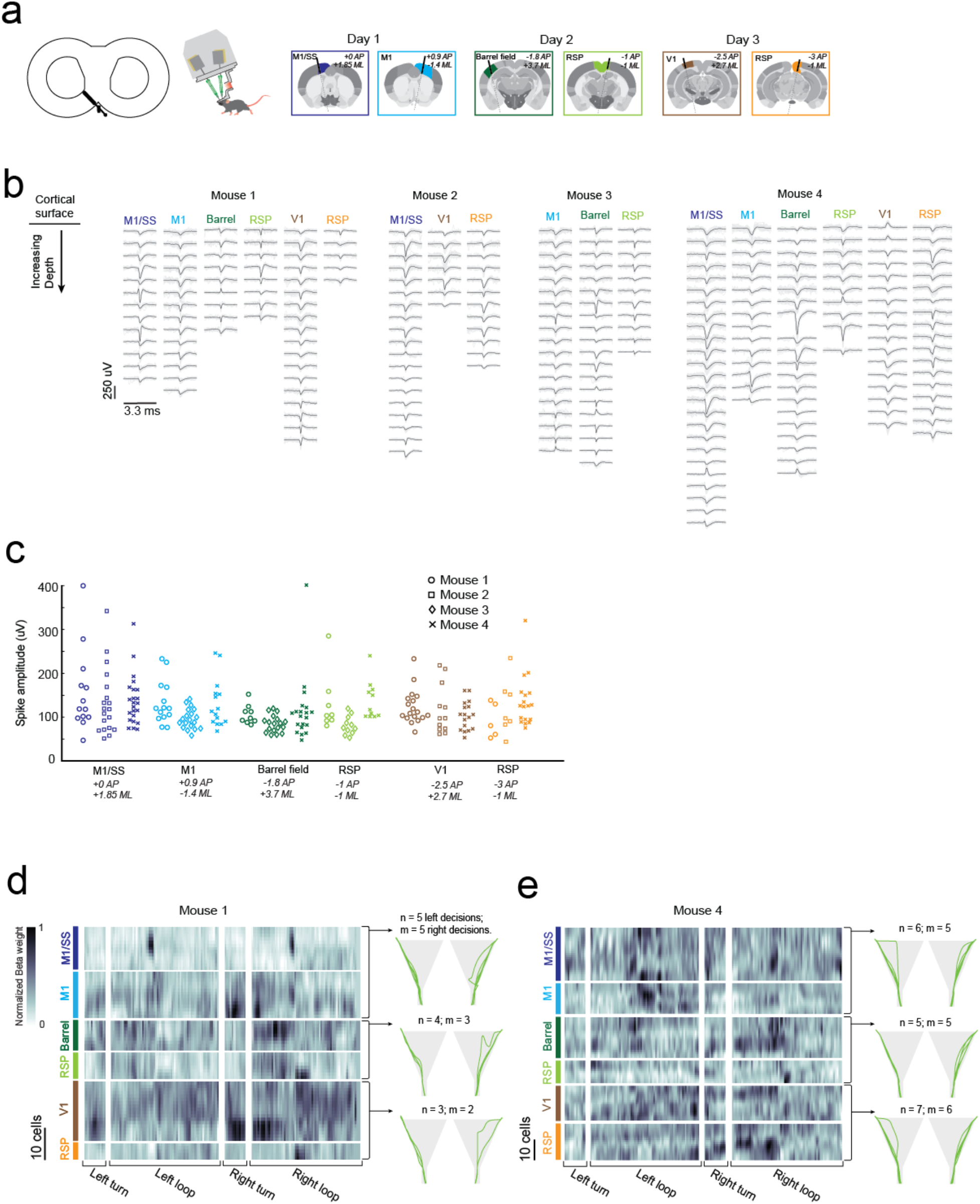
Summary of electrophysiology results across all mice (a) Symbols of the 8-maze arena with one door in place (decision-making) and mouse maneuvering the electrophysiology headstage indicating the data in the figure, and schematics showing the locations of the 6 recording sites. (b) Mean spike waveform (black) and a subsample of 50 individual spikes for each cell identified in the electrophysiology recordings in each mouse during the navigational decision-making task. (c) Spike amplitudes of each of the mean waveforms shown in (b). (d) Kernel matrices of normalized Beta weights for mouse 1, obtained using linear regression on the z-scored spike rate of each cell and the location of the mouse in the 8-maze arena (196 location bins). Kernel matrices from all 3 days are concatenated vertically with (left) the recording site labelled and color-coded, and (right) global X-Y paths taken by the mouse through the turning zone for each day. (e) As in (d) but for mouse 4.

## Supplementary Information

**Supplementary Information 1:**
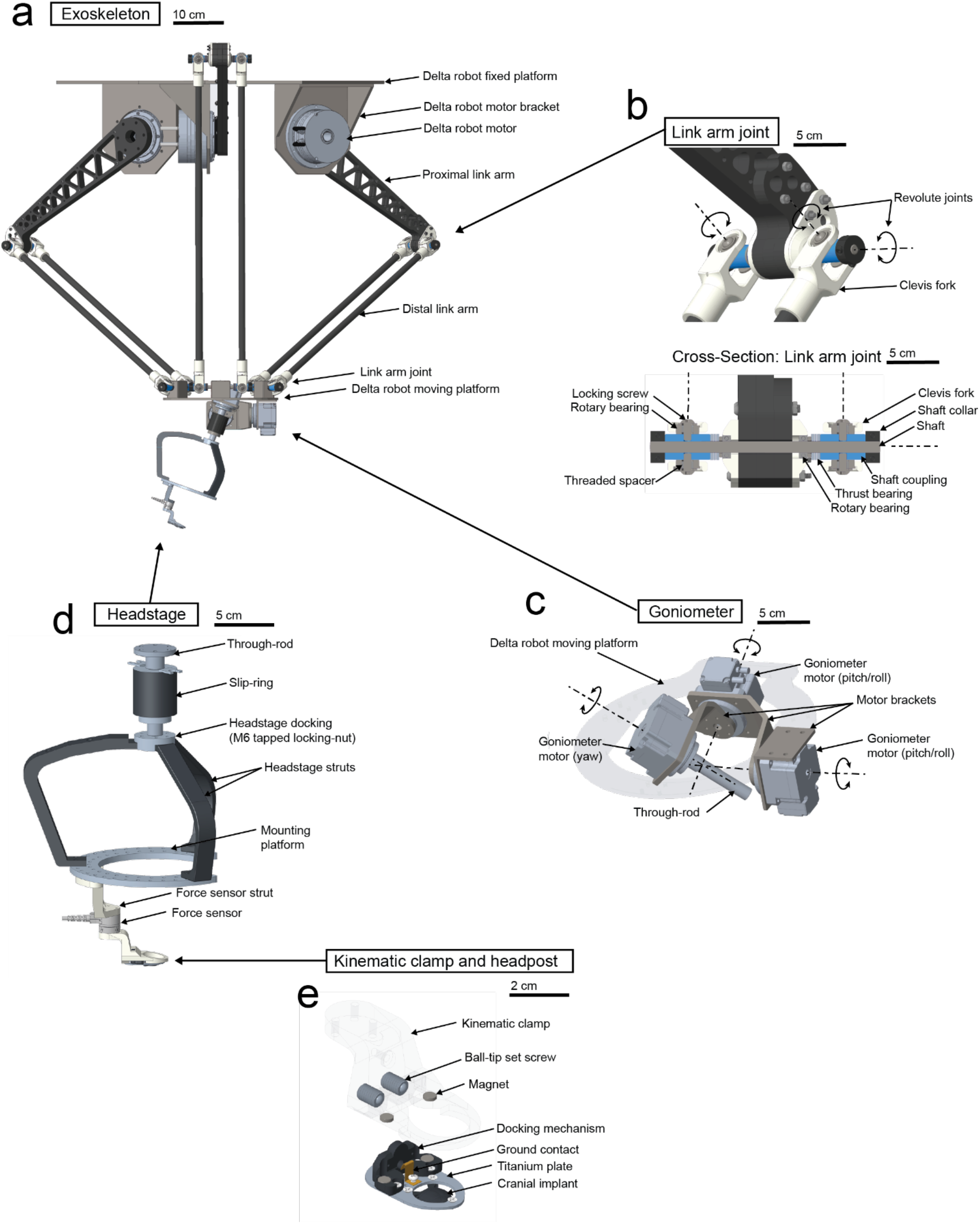
Exoskeleton construction (a) Exoskeleton delta robot, where each of the 3 motors are connected to the moving platform via a proximal arm, a pair of distal link arms, and a pair of link arm joints. (b) Delta robot link arm joint (top) and cross-section through a pair of orthogonal revolute joints showing the bearing component-stack (bottom). (c) Goniometer mounted within the moving platform of the delta robot, where 3 motors configured orthogonal to one another generate pitch/roll/yaw motion. (d) Headstage without any neural or behavioral monitoring equipment mounted to it, with the slip-ring at the top and the force sensor at the bottom. (e) Kinematic clamp and chronically implanted headpost, where ball-tip set screws and magnets facilitate docking of the mouse to the headstage.

**Supplementary Information 2:**
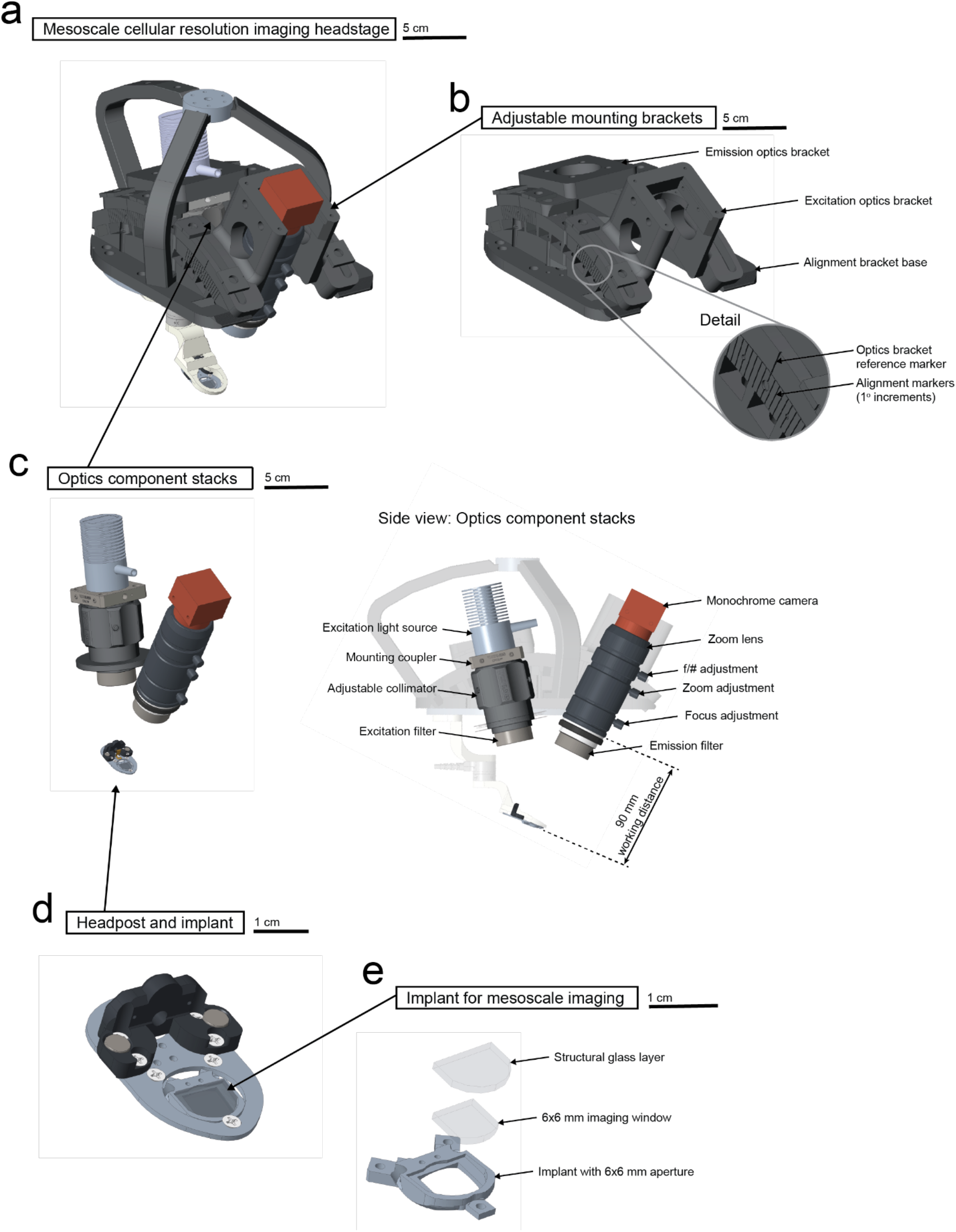
Headstage configuration for mesoscale, cellular resolution imaging (a) Headstage configuration for mesoscale imaging using 1-photon excitation. (b) Mounting brackets for the emission and excitation optics stacks can have their angle adjusted with respect to the imaging plane on the mouse’s implant. (c) Component stacks (left) for the emission and excitation optics and (right) side-on-view of these component stacks within the headstage with components and working distance annotated. (d) Headpost and implant for mesoscale imaging, that are chronically attached to the mouse. (e) Exploded view of the implant for mesoscale imaging, which is surgically implanted over the mouse’s brain, showing the 2 layers of the glass.

**Supplementary Information 3:**
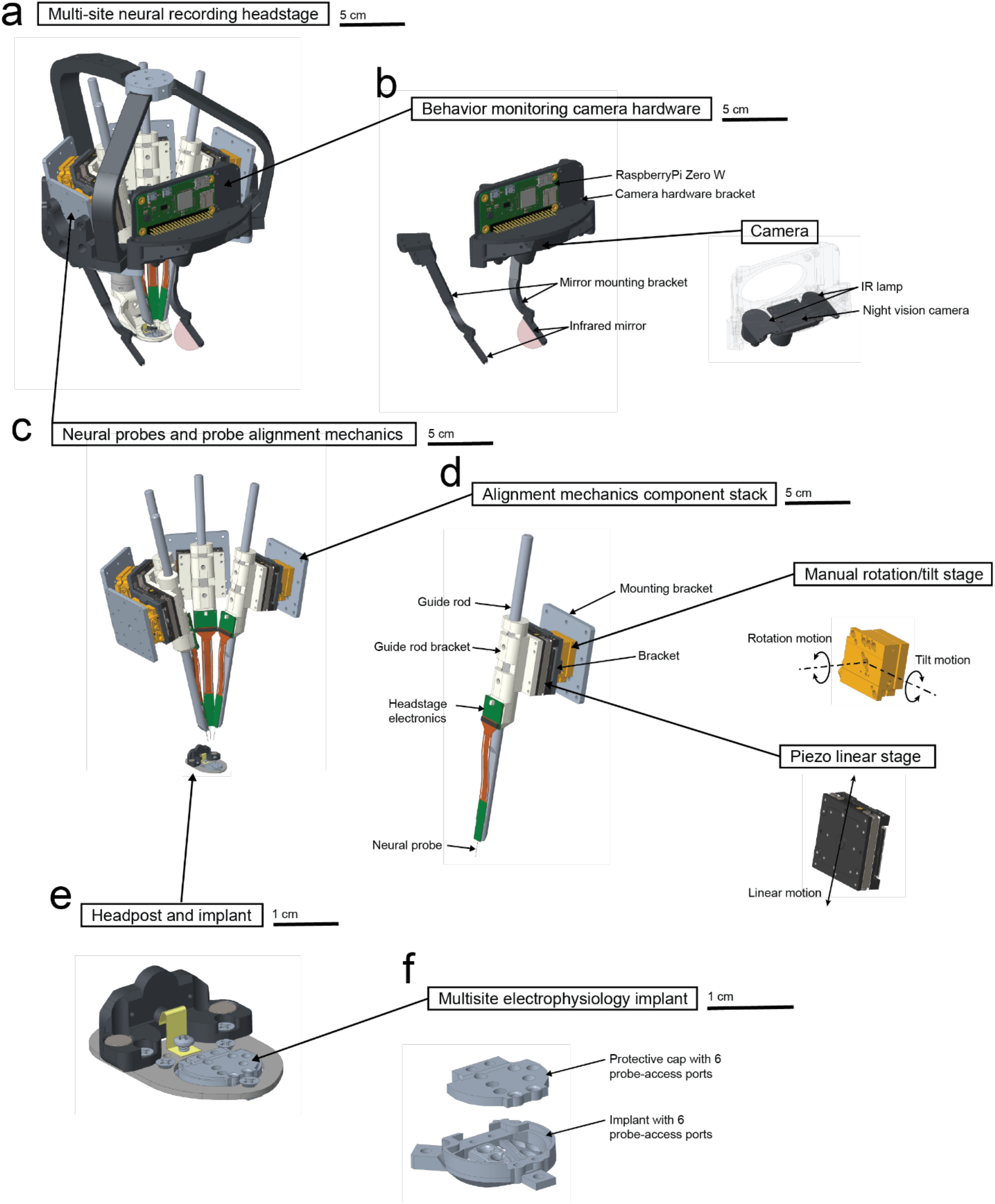
Headstage construction for multi-site electrophysiology and behavioral monitoring (a) Headstage configuration for multi-site electrophysiology and behavioral monitoring, with 4 Neuropixels probes installed. (b) Behavioral monitoring camera hardware includes (left) a microcontroller to capture and store video data, optional infrared mirrors to monitor pupil and whisker activity, and (right) a night vision camera with infrared lamps to illuminate the field of view. (c) Four sets of neural probes and probe alignment mechanics arranged above the headpost. (d) One set of alignment mechanics and neural probe with components labelled, and the axes of travel of the (top-right) manual rotation/tilt stage used to align the neural probe and the (bottom-right) piezo linear stage used to insert and remove the neural probe. (e) Headpost and implant for multi-site electrophysiology that are chronically attached to the mouse (for surgical procedure, see **Supplementary Info. 8**). (f) Exploded view of the implant for multi-site electrophysiology, where silicon is inserted between the implant and the protective cap to protect the exposed brain within the probe access-ports.

**Supplementary Information 4:**
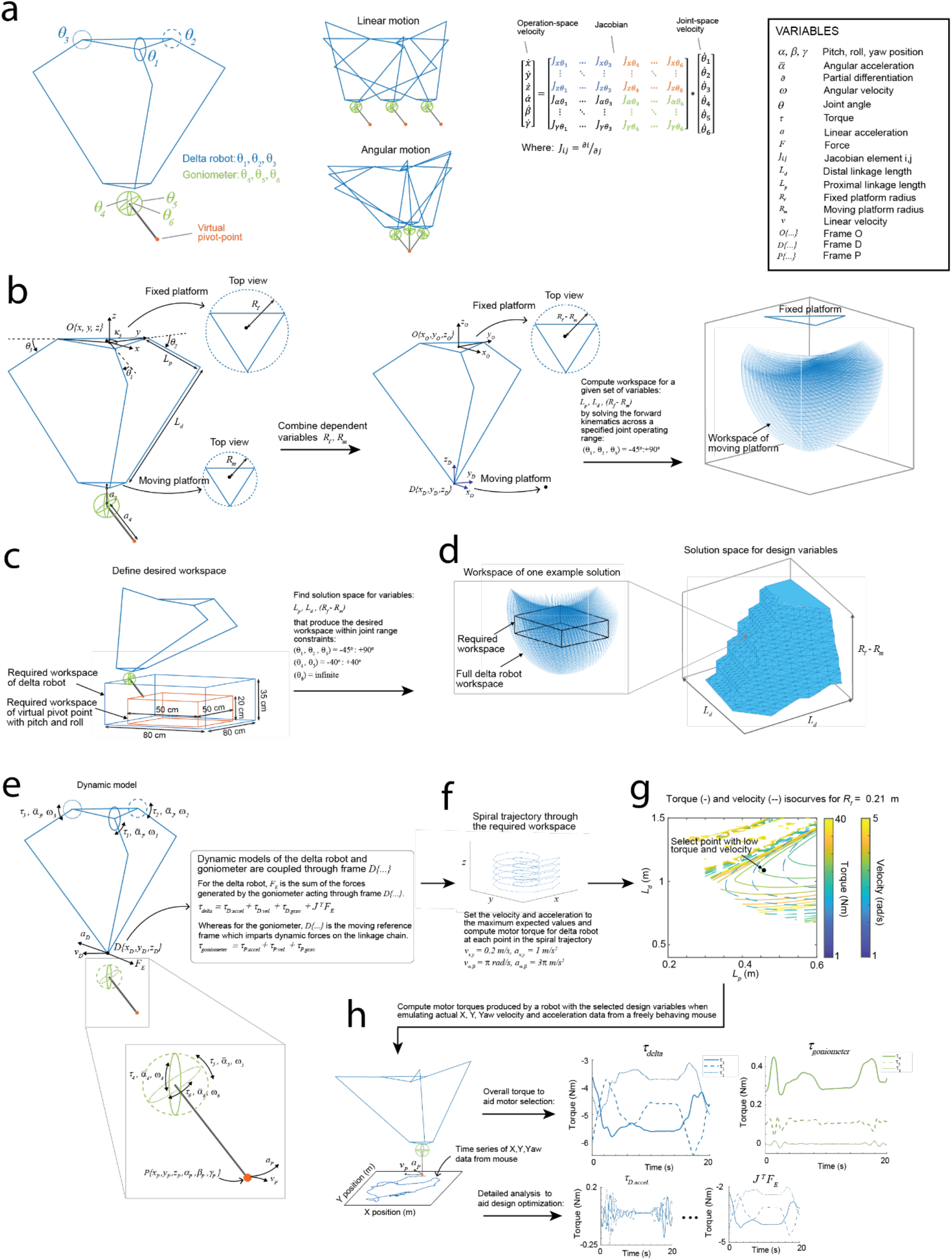
Exoskeleton design optimization (a) Schematics showing the 3 actuated joints in the delta robot and the 3 in the goniometer and motion sequences showing the synchronized movement of these joints during linear and angular motion of the pivot point. The system Jacobian with colors indicating the elements associated with linear (blue), angular (green), and pivoting (orange) motion. (b) The main design variables for the delta robot (left), combining the fixed and moving platform radii into one variable (center), and delta robot workspace (right). (c) The desired workspace (range of motion) for the delta robot and its relationship to the desired workspace for the pivot point. (d) The solution space (combinations of dimensions which can achieve the desired range of motion in the delta robot), with a schematic of one example solution (left). (e) The main variables in the dynamic model of the exoskeleton. (f) The spiral trajectory through the workspace and velocity and acceleration parameters set at each point along the trajectory. (g) The maximum motor torque and joint velocity computed along the spiral trajectory for each of the combinations of dimensions in the solution space in (d), and a point selected with both low torque and low joint velocity. (h) Time series data acquired from markerless tracking of a freely behaving mouse (left) and the corresponding time series of motor torque acceleration, velocity, gravity, and external force components (right).

**Supplementary Information 5:**
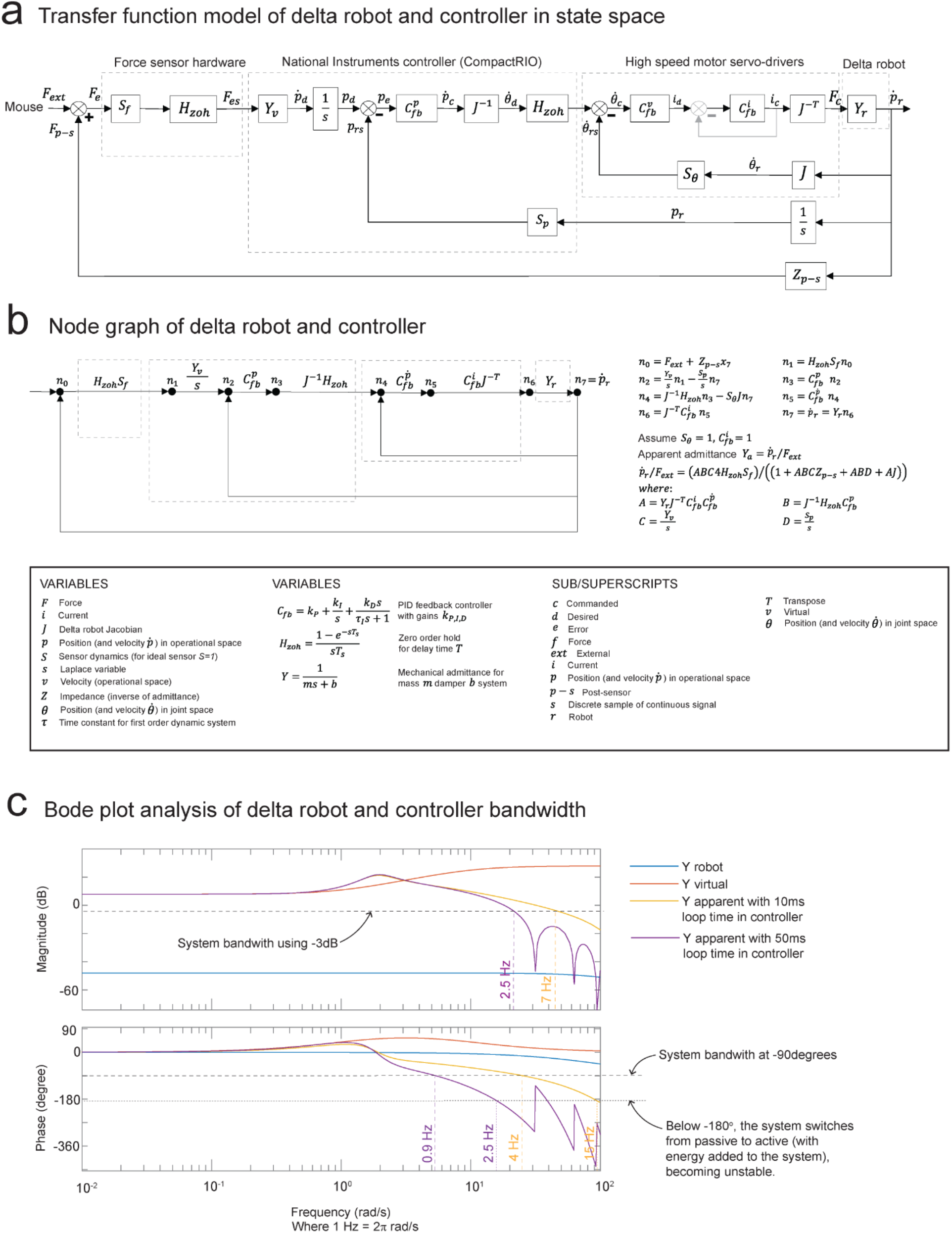
Evaluation of the system bandwidth using a Laplace space model (a) A transfer function model of the system (robotic exoskeleton and controller), with variables transformed to Laplace-space. (b) Node graph of the transfer function model in (a). (c) Bode plot (top, magnitude vs. frequency; bottom, phase vs. frequency) of the system showing the apparent admittance (yellow and purple) emulates the virtual admittance (orange) well at low frequencies but at higher frequencies degrades and approaches the true robot admittance (blue). Reducing the loop time on the controller from 50 ms (yellow) to 10 ms (yellow) increases the bandwidth from 0.9 – 2.5 Hz to 4 – 7 Hz.

**Supplementary Information 6:**
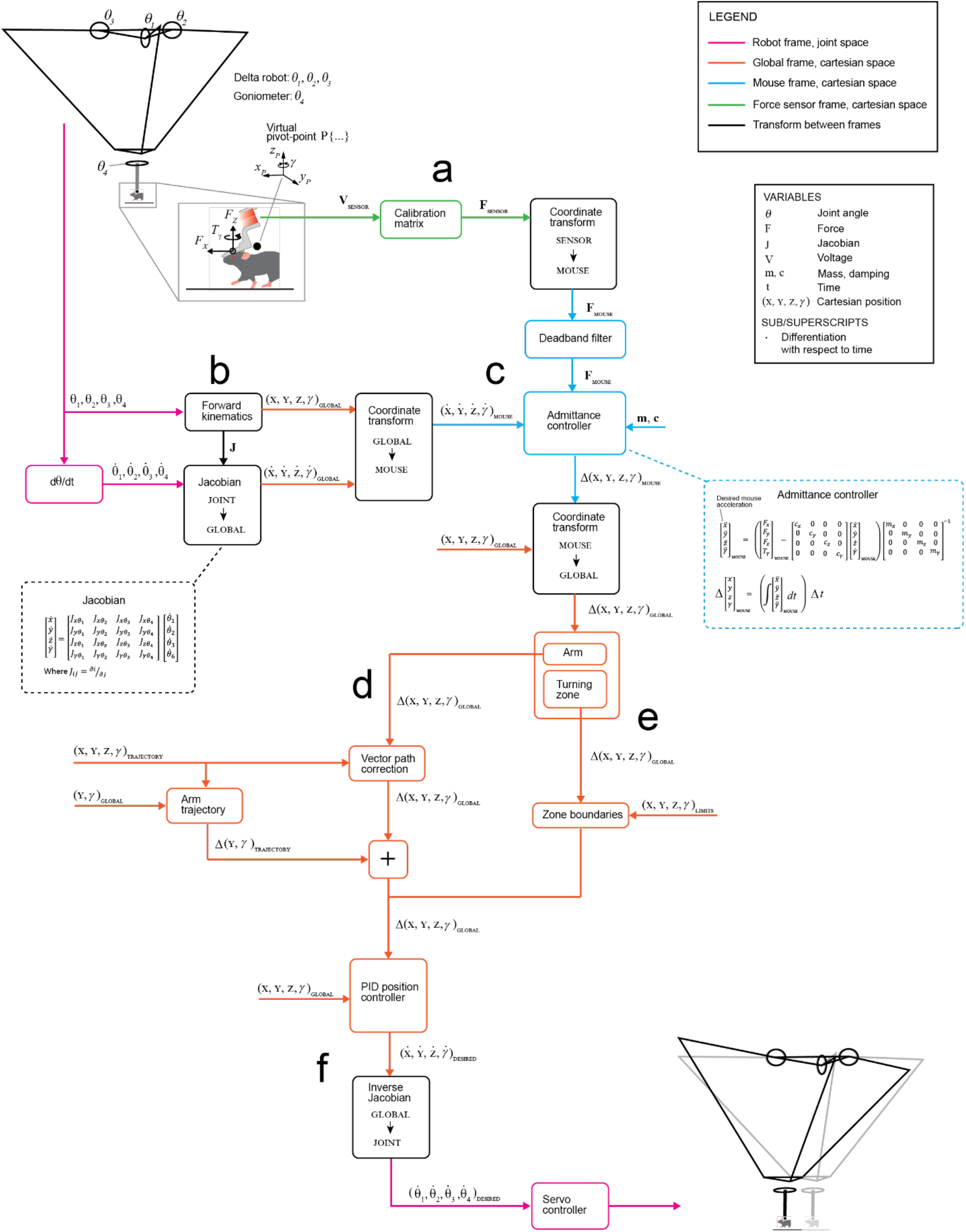
Flow diagram of the exoskeleton controller in the 8-maze arena (a) Raw voltages from the force sensor are converted to forces in the force sensor frame using a calibration matrix, and then these forces transformed to the mouse’s coordinate frame to describe the mouse’s forces. (b) Forward kinematics converts the joint angles of the robot into the global position of the mouse and to define the Jacobian. The Jacobian converts the joint velocities to global velocities which are then transformed to the mouse’s coordinate frame to describe the mouse’s velocities. (c) The mouse’s forces and velocities are fed into the admittance controller, along with programmed mass and damping values, to find the desired mouse accelerations. These accelerations are then integrated to find a desired change in position (the product of the desired velocity and controller loop time), which is transformed to the global coordinate frame. (d) If the mouse is an arm of the 8-maze arena, then the desired change in position is modified using vector path correction and the mouse’s Y and Yaw axes are trajectory controlled before being sent to the PID controller (see also **Extended Data 5**). (e) If the mouse is in the turning zone of the 8-maze arena, then the desired change in position is checked to ensure it is within the zone boundaries before being sent to the PID controller. (f) The output of the PID controller is desired velocities, which are transformed to joint space using the inverse Jacobian to produce desired joint velocities. These joint velocities are sent to the servo controllers that drive each motor in the robot to produce motion.

**Supplementary Information 7:**
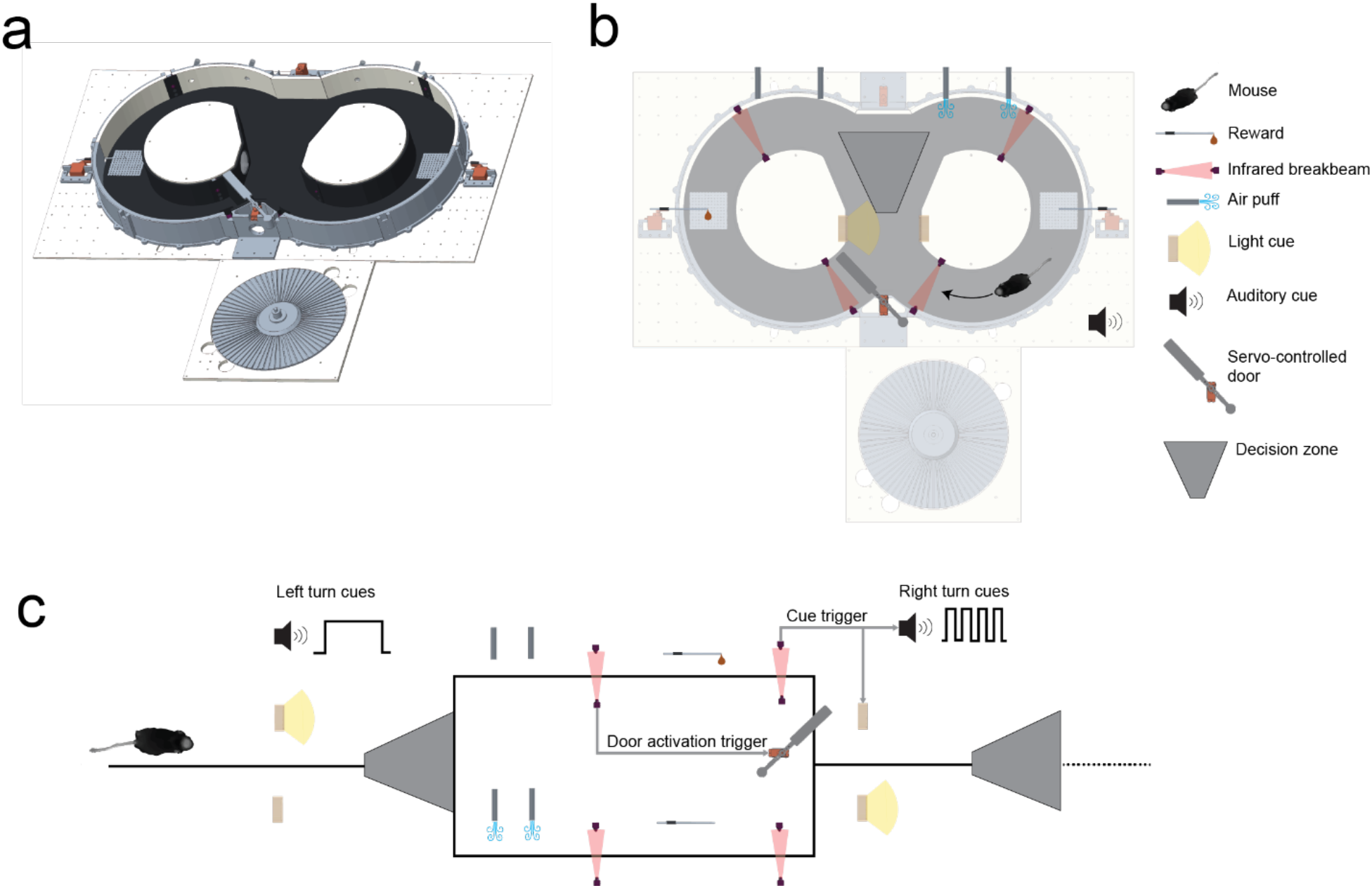
Navigational decision-making task protocol in the 8-maze arena (a) CAD rendering of the 8-maze arena. (b) Top-down view of the 8-maze arena and its elements, with the mouse in the right arm and elements configured for a left turn (right arm air-puff; left arm reward; left turn light cue; servo door turned left). (c) Linear sequence showing left turn configuration of the 8-maze elements, and the trigger operations of the break-beam sensors.

**Supplementary Information 8:**
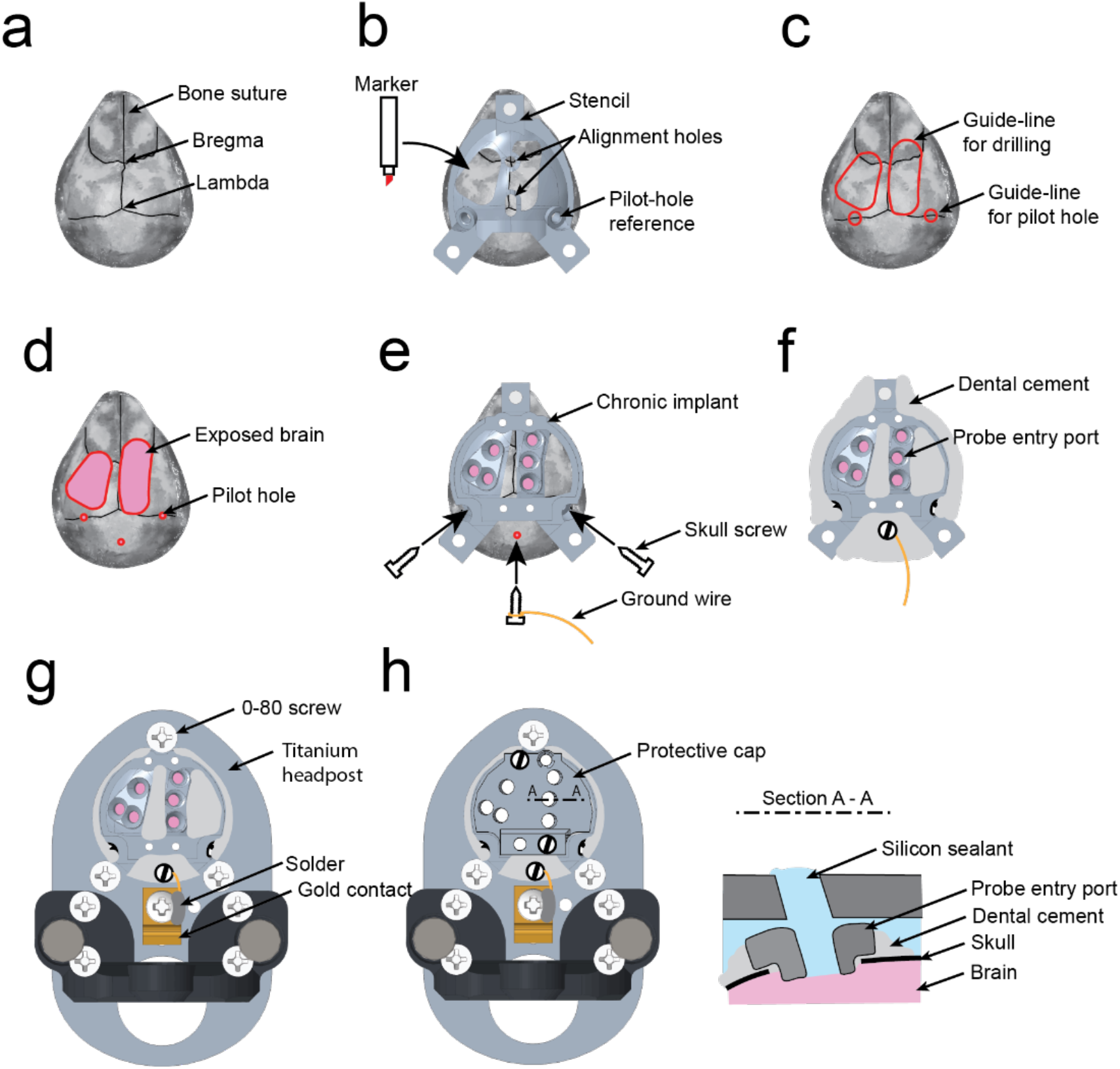
Surgical procedure for chronical implantation of the headpost and implant for multi-site electrophysiology (a) Bone sutures and anatomical landmarks visible on the dorsal surface of the skull after removing the scalp and soft connective tissue. (b) Plastic stencil used to reference the locations of the targeted craniotomies and pilot-holes for skull screws to the locations of the anatomical landmarks on the skull surface. (c) Dorsal surface of the skull after marking guide-line locations of the targeted craniotomies and pilot-holes for skull screws using the stencil in (b). (d) Areas of exposed brain tissue after craniotomy and pilot holes for skull screws. (e) Implant for multi-site electrophysiology is screwed down using 2 skull screws through 2 of the pilot holes, and a third skull screw with a ground wire attached is screwed down through the third pilot hole. (f) The implant and skull screws are secured using dental cement. (g) The headpost for electrophysiology is attached to the implant using three 0-80 screws, and the ground wire is soldered onto a gold contact located on the headpost. (h) A protective cap is screwed on to the implant using self-tapping screws (skull screws) after applying silicon sealant over the exposed brain. Before recording sessions, this silicon sealant is replaced with a silicone-gel.

## GitHub repo

### CAD

a. Exoskeleton construction
b. Electrophysiology headstage construction
c. Imaging headstage construction
d. Linear oval track arena
e. 8-maze arena

### Software

a. LABVIEW files for exoskeleton control on the CompactRIO
b. LABVIEW files for 8-maze control on the CompactRIO (freely behaving training)

### Data analysis

MATLAB files and raw data for:

a. Exoskeleton design optimization
b. Exoskeleton admittance controller tuning
c. Gait analysis
d. 8-maze task performance analysis
e. Mesoscale imaging data analysis during 8-maze navigation
f. Electrophysiology data analysis during 8-maze alternating choice task

